# Matrix mechanics gate the expression of spontaneous polarity in multicellular aggregates

**DOI:** 10.64898/2026.11.15.766081

**Authors:** Theodora Rogkoti, Apeksha Shapeti, Jorge Barrasa-Fano, Silvia Hervas-Raluy, Danail Stoychev, Giuseppe Ciccone, Graham J. Day, Oana Dobre, Manlio Tassieri, Adrian Ranga, Wolfgang A. Wall, Matthew J. Dalby, Catherine Berry, Jose Manuel García-Aznar, Marco Cantini, Hans Van Oosterwyck, Manuel Salmeron-Sanchez

## Abstract

Multicellular aggregates are initially spherical, yet tissues routinely break this symmetry to form protrusions, invade, and migrate. Whether this symmetry breaking originates within the aggregate or is imposed by its surroundings remains unclear. Here, we embed human mesenchymal stem cell (hMSC) spheroids in synthetic poly(ethylene glycol) hydrogels in which stiffness, viscoelasticity, and degradability are controlled independently. Spheroids in elastic matrices remain spherical; viscoelasticity permits symmetry breaking only in compliant networks, whereas proteolytic degradability permits it in both compliant and rigid matrices. Brillouin microscopy shows that symmetry breaking is accompanied by softening of the spheroid core and loss of internal radial symmetry. Three-dimensional traction force microscopy reveals that the force field is polarised even in spheroids that remain spherical: tractions concentrate at two approximately opposing poles, and focal adhesions adopt a similar bipolar distribution. In spheroids that do break symmetry, protrusions emerge opposite the pole of highest traction. A continuum model shows that an anisotropic distribution of contractile cells is sufficient to reproduce both the patterns and magnitude of the measured displacements. Force polarity is therefore an intrinsic property of the aggregate, whereas matrix mechanics act as a filter that determines whether this polarity is expressed as symmetry breaking.

## Introduction

Living cells reside within a complex, three-dimensional (3D) extracellular matrix (ECM) composed of fibrous proteins and polysaccharides that provide them with structural, biochemical, and mechanical cues^1^. Through adhesion receptors such as integrins, cells physically anchor to the ECM and actively probe its mechanical properties, generating and responding to forces that are transmitted across multiple length scales^2^. These reciprocal mechanical interactions allow the ECM to function as an information channel, conveying mechanical cues that guide cellular responses through a process known as mechanotransduction.

Over the past decades, the mechanical properties of the ECM have emerged as central regulators of cell behaviour and function in various physiological and pathological processes, including morphogenesis^3^, wound healing and fibrosis^4^, and cancer progression^5,6^. Consequently, understanding how cells sense and respond to ECM mechanics has become a foundational question in mechanobiology. Early studies largely focused on tissue elasticity, often quantified as stiffness, as a primary mechanical determinant of cell behaviour. These works demonstrated that matrix stiffness can regulate cell spreading^7^, migration^8,9^, proliferation^9^, and lineage specification^10,11^ in both two-dimensional (2D) and 3D hydrogel systems, establishing elasticity as a powerful design parameter for bioengineered in vitro models. While these insights have profoundly shaped our understanding of mechanotransduction and inspired a widespread use of tuneable elastic hydrogels, they do not capture the mechanical complexity of native tissues.

In vivo, tissues are not purely elastic but instead exhibit viscoelastic behaviour, characterised by time-dependent stress relaxation and energy dissipation^12^, and are continuously remodelled through cell-mediated degradation^13^. These properties arise from the dynamic reorganisation and breakdown of ECM networks, respectively, and are increasingly recognised as fundamental features of the cellular microenvironment^12,14^. Recent studies using cell-laden 3D hydrogel systems have shown that stress relaxation can profoundly influence single-cell behaviours such as spreading^15,16^, migration^17^, and differentiation^18^, often eliciting responses comparable to those previously observed in degradable 3D matrices^7^. However, viscoelasticity and degradability are usually coupled within a single material system, or studied separately in different systems, so their distinct mechanoregulatory roles remain difficult to separate. Moreover, most studies have examined their effects at the single-cell level, leaving open questions about how they regulate collective cell behaviour.

This matters, because cells in vivo rarely act in isolation. Instead, they function as multicellular collectives during processes such as tissue development, regeneration, and disease progression, where mechanical forces are coordinated across many cells and transmitted through both cell-cell and cell-matrix interactions^19,20^. Yet collective responses remain poorly modelled in 3D environments with physiologically relevant mechanical properties. Multicellular spheroids provide a powerful in vitro model for studying such collective behaviours as they recapitulate key features of native tissues, including cell–cell interactions, emergent mechanical properties, and coordinated force generation^21^. MSC spheroids, in particular, have attracted interest due to their enhanced regenerative potential, improved survival, and pro-healing secretory profiles compared to single cells^22,23^. When embedded within hydrogels, MSC spheroids can actively remodel their surroundings and promote vascularisation and tissue repair, making them promising building blocks for tissue engineering and regenerative therapies^24^. Previous studies have demonstrated that hydrogel properties, such as stiffness and ligand presentation, can influence spheroid spreading and differentiation^25^, processes important for their regenerative capacity. However, how viscoelasticity and degradability regulate spheroid-level properties such as stiffness, and contractility remains largely unknown. More fundamentally, spheroid spreading requires an initial spherical aggregate to break symmetry^26,27^, and it is unclear whether the direction of this symmetry breaking is encoded within the aggregate or imposed by the surrounding matrix.

To separate the contributions of matrix stiffness, viscoelasticity, and degradability to collective cell behaviour, we embed hMSC spheroids in three-dimensional ECM-mimetic poly(ethylene glycol) (PEG) hydrogels in which each of these properties is controlled independently. We combine Brillouin microscopy and 3D traction force microscopy (TFM) to quantify passive and active cell mechanics, respectively, while monitoring focal adhesion (FA) organisation and protrusion formation. These measurements are complemented with a simplified computational model showing that an anisotropic distribution of contractile cells within the spheroid is sufficient to reproduce both the spatial pattern and magnitude of the measured matrix displacements. Together, our findings demonstrate that force polarity is an intrinsic property of the aggregate and that matrix mechanics act as a filter that determines whether this polarity translates into symmetry breaking.

## Results

### Engineered viscoelasticity and degradability in fibronectin (FN)–functionalised PEG hydrogels

Two distinct groups (soft, S; and rigid, R) of mechanically tunable, synthetic hydrogels (elastic, E; viscoelastic, V; and degradable, D) were produced using eight-arm PEG-maleimide (8PEG-MAL) macromers crosslinked with linear PEG-dithiol (HS-PEG-SH) polymers or peptides and biologically functionalised with full-length PEGylated fibronectin (PEG-FN)^28^ to preserve its multidomain structure. The stiffness was raised by increasing the 8PEG-MAL weight fraction, while viscoelasticity was accomplished by varying the molecular weight of PEG-MAL polymers and PEG-dithiol crosslinkers, an approach previously established in a range of systems^29,30^. Increasing polymer chain length was expected to enhance molecular flexibility and promote physical entanglements, thereby enhancing the dissipative properties and stress relaxation rates of these networks. Shorter, less flexible chains were expected to yield more elastic, energy-retaining networks. Unlike previously reported viscoelastic systems, in which energy dissipation arises from the dissociation of weak dynamic bonds^31^, viscoelasticity in this system emerges from dissipative motion and rearrangement of flexible polymer chains within the aqueous, covalently crosslinked hydrogel network. Finally, controlled degradability was introduced into the elastic formulations by incorporating a matrix metalloproteinase (MMP)-sensitive peptide crosslinker (VPM; GCRDVPMSMRGGDRCG)^28^ at a 1∶1 molar ratio with PEG-dithiol to generate networks susceptible to enzymatic cleavage (**Fig. 1a,b** and **Table S1**).

**Figure 1:**
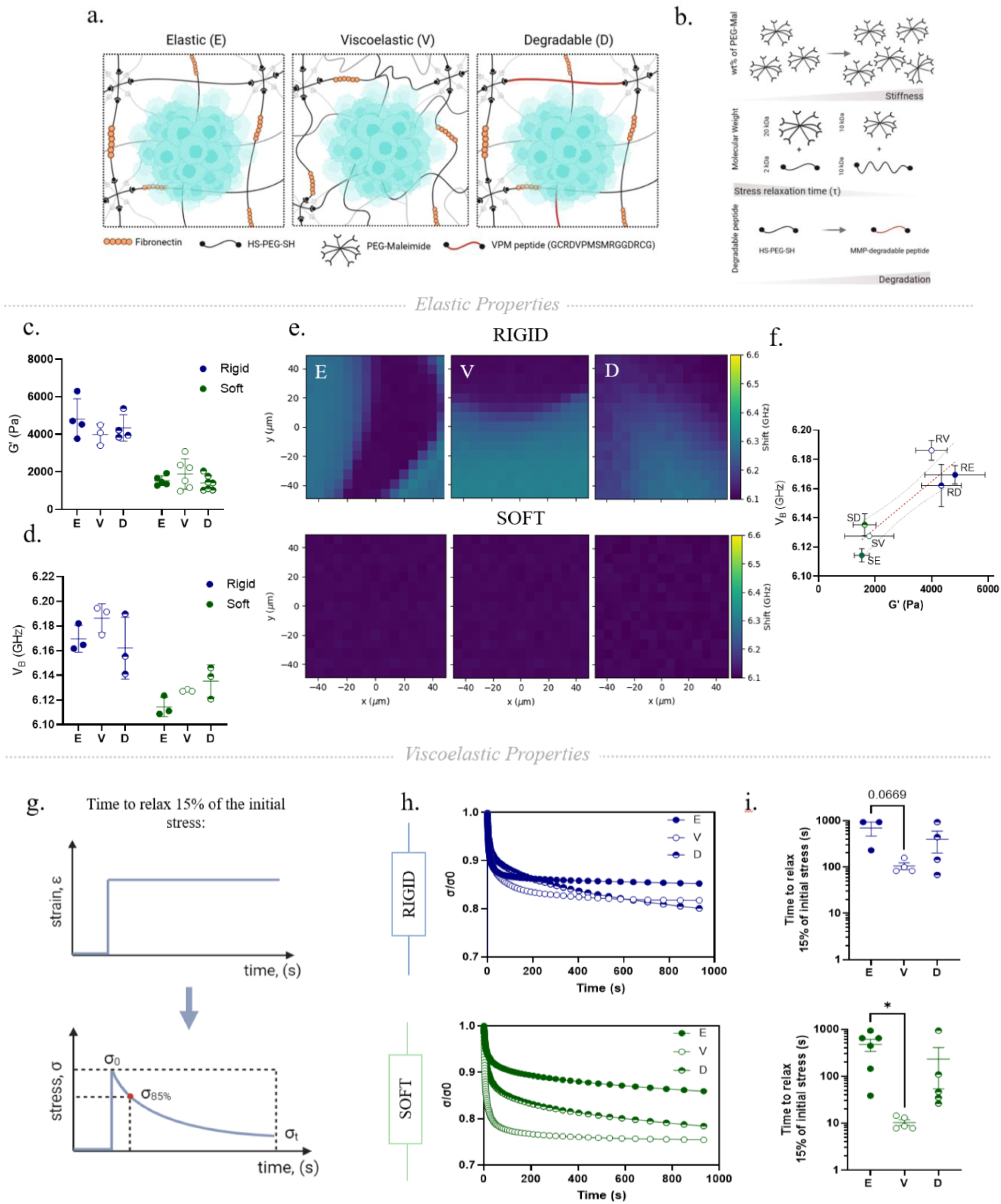
Mechanical characterisation of PEG-MAL hydrogels. **(a)** Schematic representation of different hydrogel networks. Image created using BioRender. **(b)** Approaches used to modify hydrogel stiffness, viscoelasticity, and degradability in PEG-MAL hydrogels. Image created using BioRender. **(c)** Shear rheology measurements, showing storage modulus (*G’*) data obtained from rigid (4.5 kPa, blue) and soft (1.5 kPa, green) hydrogel families. Data are shown as mean ± SD, RE: n = 4, RV: n = 3, RD: n = 4 and SE: n = 5, SV: n = 6, SD: n = 6 hydrogels. One-way ANOVA with Tukey’s post-hoc test for multiple comparisons. **(d)** Quantification of the average V_B_ for each hydrogel condition. Data are shown as mean ± SD, n = 3 hydrogels per condition. Two-way ANOVA with Tukey’s post-hoc test for multiple comparisons. **(e)** Representative Brillouin frequency shift (V_B_) maps of rigid (top row) and soft (bottom row) elastic, viscoelastic and degradable hydrogels. **(f)** Average V_B_ values plotted against the storage modulus (*G’*) for all hydrogels. Simple linear regression analysis, R^2^ = 0.6628. Dotted black lines represent the 95% confidence intervals of the best-fit line (dotted red line). **(g)** Graphical representation of stress relaxation test. During stress relaxation tests, the material experiences a constant strain (ε) over a prolonged period of time (t). In viscoelastic materials, the resulting stress-relaxation curve (blue) will show a gradual decrease of stress (σ) over time. The time it takes to relax 15% of the initial stress (σ_85%_ of stress remaining) is used here as a parameter to assess the differences in time-dependent mechanical properties between different hydrogels. **(h)** Representative average stress relaxation curves are shown for rigid and soft hydrogel families, where the stress (σ) is normalised to the maximum initial stress (σ0). 1% and 10% strain was applied for rigid and soft hydrogels, respectively. **(i)** The time it takes to relax 15% of the initial stress is calculated for rigid and soft hydrogel families. Data is shown as mean ± SD, RE: n = 3, RV: n = 4, RD: n = 4 and SE: n = 6, SV: n = 5, SD: n = 5 hydrogels. *\*p-value* < 0.05, One-way ANOVA with Tukey’s post-hoc test for multiple comparisons. No annotations signify no statistical significance.

Shear rheological measurements of bulk hydrogels revealed no significant differences between the elastic moduli (*Gʹ*) of SE, SV, and SD (*Gʹ* ≈ 1.5 kPa) and RE, RV, and RD (*Gʹ* ≈ 4.5 kPa), confirming the hydrogels were isoelastic within each stiffness group (**Fig. 1c and Supplementary Fig. 1a-d**). To further assess the mechanical properties at the microscale, we additionally employed Brillouin spectroscopy, a non-contact optical technique that provides high-resolution, label-free mechanical mapping through Brillouin light scattering (BLS). BLS generates a quantifiable Brillouin frequency shift (V_B_), which is directly related to the complex, frequency-dependent longitudinal modulus (*M*) and serves as a proxy for local stiffness; higher V_B_ values correspond to stiffer materials, whereas lower V_B_ values correspond to softer materials or liquids^32^. Spatially resolved Brillouin frequency shift maps revealed comparable V_B_ values among hydrogels within each stiffness group (**Fig. 1d,e**; soft: V_B_ ≈ 6.13 GHz; rigid: V_B_ ≈ 6.17 GHz), which linearly correlated with the *Gʹ* measurements, confirming the isoelastic nature of the two hydrogel groups at both the bulk-scale and microscale (**Fig. 1f**). Finally, SV and RV hydrogels displayed greater viscoelastic character than their elastic counterparts, with faster times to relax 15% of an applied constant stress (**Fig. 1g,h**), indicating an enhanced capacity for energy dissipation within these hydrogels (**Fig. 1i**).

Stable long-term retention of PEG-FN within the network was achieved by FN PEGylation which allowed the integration of the protein in the hydrogel backbone and promoted direct cell engagement with the bulk material rather than with a loosely adsorbed protein^28^. Successful incorporation of PEGylated FN was confirmed by imaging fluorescently tagged PEG-FN, which revealed no significant differences in FN content between elastic, viscoelastic, and degradable hydrogels in either stiffness family (**Supplementary Fig. 2a-c**).

Together, these results establish an FN-based hydrogel system in which viscoelasticity and degradation can be tuned independently of the elastic modulus across different hydrogel stiffnesses, making it a well-suited platform for investigating MSC multicellular responses to distinct ECM mechanical cues.

### Matrix mechanics determine whether spheroids break shape symmetry

Previous studies have shown that viscoelastic and degradable hydrogels can each promote the spreading of single MSCs in 3D environments^33,34^, but these studies have typically examined a single stiffness regime and have not addressed multicellular aggregates. To investigate which mechanical features of the matrix permit a spheroid to break shape symmetry, small hMSC spheroids (approximately 100 cells per spheroid) were encapsulated in soft (SE, SV, and SD) and rigid (RE, RV, and RD) hydrogels.

Viability assessment after five days of culture revealed that hMSC spheroids remained viable across all conditions, with only a few dead cells present in spheroid cores, indicating that the hydrogels did not compromise cell survival (**Supplementary Fig. 3a,b**). These findings are consistent with previous studies demonstrating that small spheroids do not develop pronounced hypoxic cores^35^, supporting the use of this spheroid size to minimise hypoxia-associated cell death.

Spheroid morphology was then monitored over five days of culture. In rigid hydrogels, spheroids encapsulated in RE and RV hydrogels retained compact, spherical morphologies throughout the five-day culture period (**Fig. 2a**). Quantitative analysis of fluorescent confocal images of the spheroids revealed minimal changes in spheroid area and circularity over time, with no significant differences observed between elastic and viscoelastic conditions (**Fig. 2b**). In contrast, when the cells were able to degrade the hydrogels, spheroids exhibited pronounced spreading, characterised by the emergence of protrusions (**Fig. 2a**). There was a clear time-dependent increase in spreading (**Fig. 2b, Table S2**), with spheroid areas significantly increasing in RD hydrogels compared to those in non-degradable RE or RV matrices by day five. Circularity measurements further supported this protrusive phenotype, with degradable matrices exhibiting significantly reduced circularity at all time points, and a progressive decline over time (**Fig. 2c, Table S2**). Together, these findings suggest that in rigid hydrogels, viscoelastic relaxation alone was insufficient to overcome the physical constraints imposed by a rigid, non-degradable network, so spheroid spreading was restricted. Only when the matrix was degradable could cells break down and reorganise the surrounding network, enabling extensive spreading and loss of spheroid symmetry.

**Figure 2:**
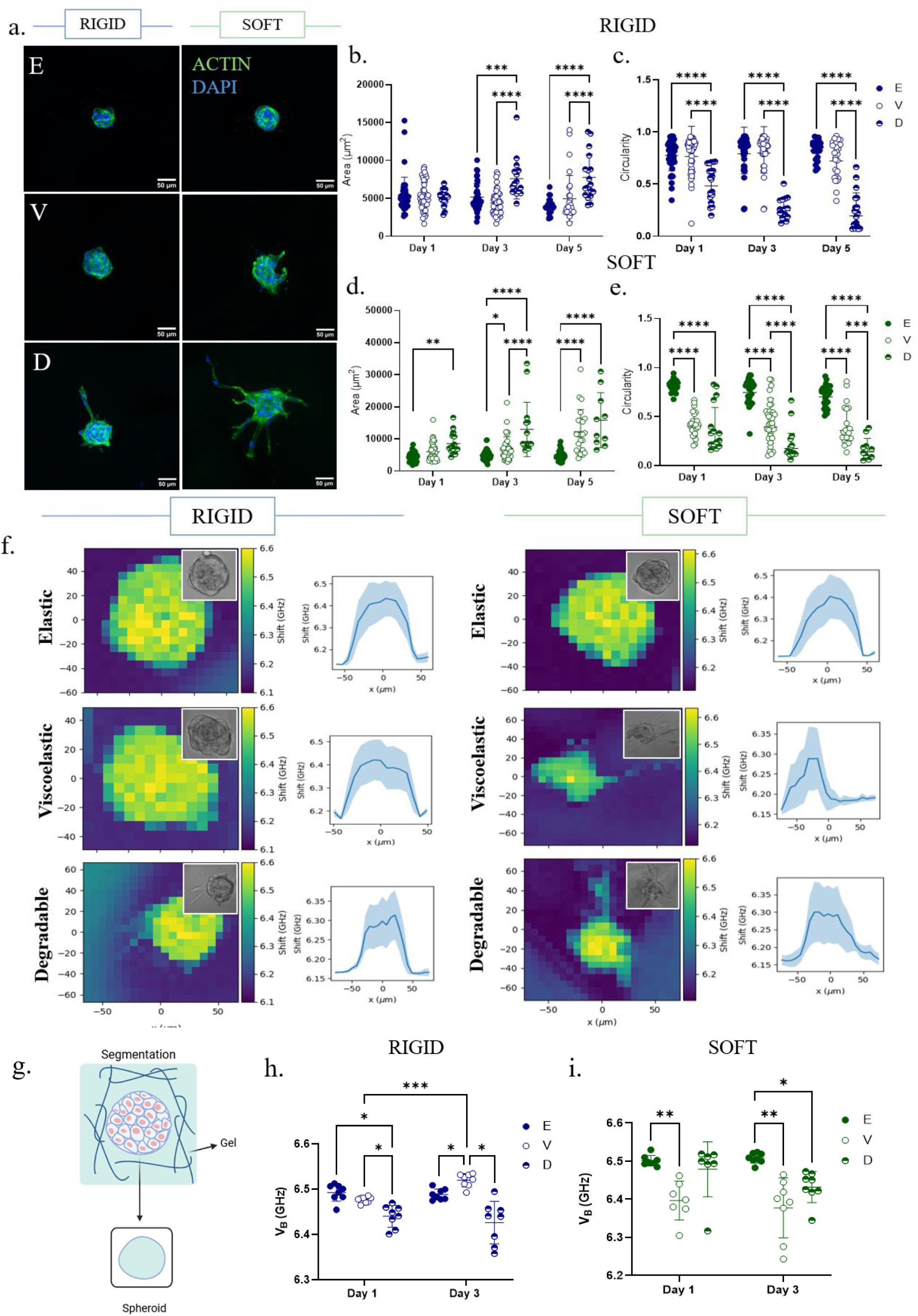
Spheroid morphological and mechanical responses to matrix viscoelasticity and degradation. **(a)** Representative immunofluorescence images of hMSC spheroids (actin cytoskeleton = green, DAPI = blue) encapsulated in different hydrogels after 5 days. Scale bar = 50 μm. **(b-c)** Quantified spheroid area and **(d-e)** circularity in rigid (top, blue) and soft (bottom, green) elastic, viscoelastic and degradable hydrogels. Data are shown as mean ± SD. RE: n = 24-44, RV: n = 29-51, RD: n = 16-19 spheroids and SE: n = 32-43, SV: n = 23- 41, SD: n = 10-15 spheroids between the two time points, N = 3. Two-way ANOVA with Tukey’s post-hoc test for multiple comparisons. *\*p-value* < 0.05, *\*\*p-value* < 0.01, *\*\*\*p-value* < 0.001, *\*\*\*\*p-value* < 0.0001. **(f)** Representative Brillouin frequency (V_B_) maps and average V_B_ line profiles of spheroids encapsulated in different hydrogel conditions on day 3. **(g)** Schematic representation of the segmentation process used to segment and analyse the VB of spheroids, excluding the surrounding microenvironment. **(h)** Quantification of the average V_B_ of spheroid cores in rigid and **(i)** soft elastic, viscoelastic and degradable hydrogels. Data is shown as mean ± SD, RE: n = 8, RV: n = 8, RD: n = 8 spheroids and SE: n = 8, SV: n = 7-8, SD: n = 7-8 spheroids between the two time points, N = 2. Two-way ANOVA with Tukey’s post-hoc test for multiple comparisons. *\*p-value* < 0.05, *\*\*p-value* < 0.01, *\*\*\*p-value* < 0.001.

In soft hydrogels, the morphology of spheroids was strongly influenced by both matrix viscoelasticity and degradability (**Fig. 2a**). Specifically, spheroids encapsulated in SE hydrogels remained largely spherical, with minimal invasive activity observed throughout the culture period, consistent with the absence of significant changes in spheroid area and circularity over time. In contrast, spheroids in SV hydrogels exhibited more protrusive and elongated morphologies, reflected by significantly increased spheroid projected areas (**Fig. 2d**) and reduced circularity (**Fig. 2e**) compared to spheroids in soft elastic hydrogels. These findings indicate that, at lower stiffness, stress relaxation facilitates collective spheroid spreading even in the absence of matrix degradation. To determine whether this spreading was associated with cell proliferation, Ki-67 immunostaining was performed. The percentage of Ki-67-positive cells was 1% in SE and 0% in SV hydrogels **(Supplementary Fig. 4a,b)**, indicating that the increased spreading in SV hydrogels was not driven by cell proliferation.

Degradation elicited the most pronounced morphological response. Spheroids encapsulated in SD hydrogels exhibited extensive spreading characterised by numerous elongated protrusions (**Fig. 2a**), leading to spheroids with significantly larger areas and reduced circularity compared to those encapsulated in either SE or SV hydrogels (**Fig. 2d-e**). A significantly higher percentage of Ki-67-positive cells was detected in SD hydrogels, indicating that proliferative activity occurred under degradable conditions and may contribute to the increased spheroid area observed in this condition **(Supplementary Fig. 4a,b)**. Similar to RD hydrogels, a time-dependent increase in spreading was also observed for both viscoelastic and degradable hydrogels at this stiffness as reflected by significant changes in both spheroid area and circularity over time **(Table S3)**.

Together, these findings demonstrate that in softer matrices, elastic networks restrict spheroid expansion, whereas both matrix viscoelasticity and degradation promote spreading to different degrees. The greater spreading observed in degradable hydrogels may reflect a higher capacity for local matrix remodelling, which is not possible through relaxation alone, as well as an increased proliferative capacity which appeared absent in non-degradable elastic or viscoelastic conditions.

Finally, a significant difference was observed in spheroid areas between rigid and soft hydrogels under degradable conditions alone (**Supplementary Fig. 5a,b**). Specifically, spheroids in SD hydrogels exhibited significantly larger projected areas and notably decreased circularities over time compared to spheroids in RD hydrogels, indicating that stiffness and hence the degree of physical confinement, sets the extent of spreading even at matched degradability.

### Spheroid symmetry breaking is associated with spheroid core softening and loss of radial mechanical symmetry

Having established that hydrogel viscoelasticity and degradability differentially regulate spheroid morphology, we asked whether these morphological changes were accompanied by changes in the mechanical phenotype of hMSC spheroids. Brillouin maps were acquired across the central plane of each spheroid, extending several micrometres into the surrounding hydrogel. From the acquired spectra, the Brillouin frequency shift (V_B_) was derived for each x-y coordinate within the mapped region. Representative Brillouin maps revealed a clear mechanical contrast between spheroids and the surrounding hydrogel, with spheroids consistently exhibiting higher Brillouin frequency shift values than their surrounding hydrogel, indicative of a higher longitudinal modulus. Across all hydrogel conditions, V_B_ values peaked at the spheroid core, as confirmed by average line profiles along the X-axis of each map (**Fig. 2f**).

Spheroids were subsequently segmented from the surrounding hydrogel region, and the average Brillouin frequency shift of the spheroid core was quantified by averaging all pixel values within the spheroid mask (**Fig. 2g, Supplementary Fig. 6**), while excluding the cell-gel interface region. Spheroids encapsulated in RE and RV hydrogels displayed similar internal stiffness on day 1, while on day 3 spheroids in RV displayed significantly elevated core stiffness compared to those in RE. In contrast, spheroids encapsulated in rigid degradable hydrogels exhibited significantly lower mean core V_B_ values compared to those in elastic and viscoelastic hydrogels at both time points, with this difference becoming more pronounced over time (**Fig. 2h**).

Both viscoelastic and degradable matrices significantly influenced the internal mechanical phenotype of hMSC spheroids in soft hydrogels. Representative Brillouin maps and corresponding average line profiles revealed marked morphological and mechanical differences among spheroids encapsulated in SE, SV and SD hydrogels. Spheroids in SV and SD hydrogels exhibited heterogeneous, asymmetric mechanical profiles with reduced core values, in contrast to the uniform, radially symmetric mechanical profiles in SE hydrogels, showing that the loss of symmetry extends to the internal mechanical field and is not confined to the outline of the aggregate (**Fig. 2f**). Quantification confirmed that the internal stiffness of spheroids in SE hydrogels did not change over three days, whereas those in SV and SD hydrogels exhibited significantly reduced Brillouin frequency shifts compared to those in SE hydrogels by day 3 (**Fig. 2i**).

Although the effect of matrix viscoelasticity on the in situ stiffness of spheroids has not, to our knowledge, been investigated, lower Brillouin frequency shifts have been reported for cancer spheroids encapsulated in degradable matrices compared with those in stiffness-matched non-degradable matrices, consistent with the trend observed here^36^. Together, these findings suggest that matrix stiffness, viscoelasticity, and degradability can differentially regulate both the morphology and internal mechanical phenotype of MSC spheroids. This may reflect the greater capacity of these matrices to accommodate cell-mediated deformation and remodelling, potentially allowing cells to extend into the surrounding matrix and altering the mechanical organisation of the spheroid.

### Cell-matrix adhesion organisation differs depending on matrix mechanics

Having established that matrix stiffness, viscoelasticity and degradability differentially regulate MSC spheroid morphology and stiffness in 3D, we next asked whether cell-matrix adhesions differ depending on matrix mechanics. Specifically, we examined the localisation and distribution of paxillin, a key focal adhesion (FA) adaptor protein recruited upon integrin engagement, in spheroids encapsulated within SE, SV, and SD hydrogels, because only at this stiffness did both viscoelasticity and degradability alter spheroid morphology.

Paxillin was detected in all conditions, irrespective of matrix mechanical properties. In elastic hydrogels, where spheroids retained a rounded morphology, overall paxillin intensity was higher than in viscoelastic hydrogels and significantly greater than in degradable hydrogels (**Fig. 3a,b**). In these elastic hydrogels, paxillin was strongly enriched at the spheroid periphery, forming bright, adhesion-rich regions, whereas in viscoelastic and degradable hydrogels, adhesion-rich areas were located at both the spheroid periphery and in cell protrusions. Quantitative analysis of FA metrics revealed that spheroids in SE hydrogels formed significantly more adhesions, which collectively occupied a greater total adhesion area than those in viscoelastic and degradable hydrogels (**Fig. 3c-e**). In contrast, the average FA areas formed by spheroids in SV hydrogels were significantly larger (although fewer in number) compared to those in elastic hydrogels (**Fig. 3f**). This difference suggests that matrix viscoelasticity may promote the formation of fewer but larger adhesions, whereas elastic matrices support the formation of a larger number of smaller adhesions. Together, these results indicate that hMSC spheroids engage the surrounding matrix through distinct FA patterns that are highly sensitive to its mechanical properties (**Fig. 3g**).

**Figure 3:**
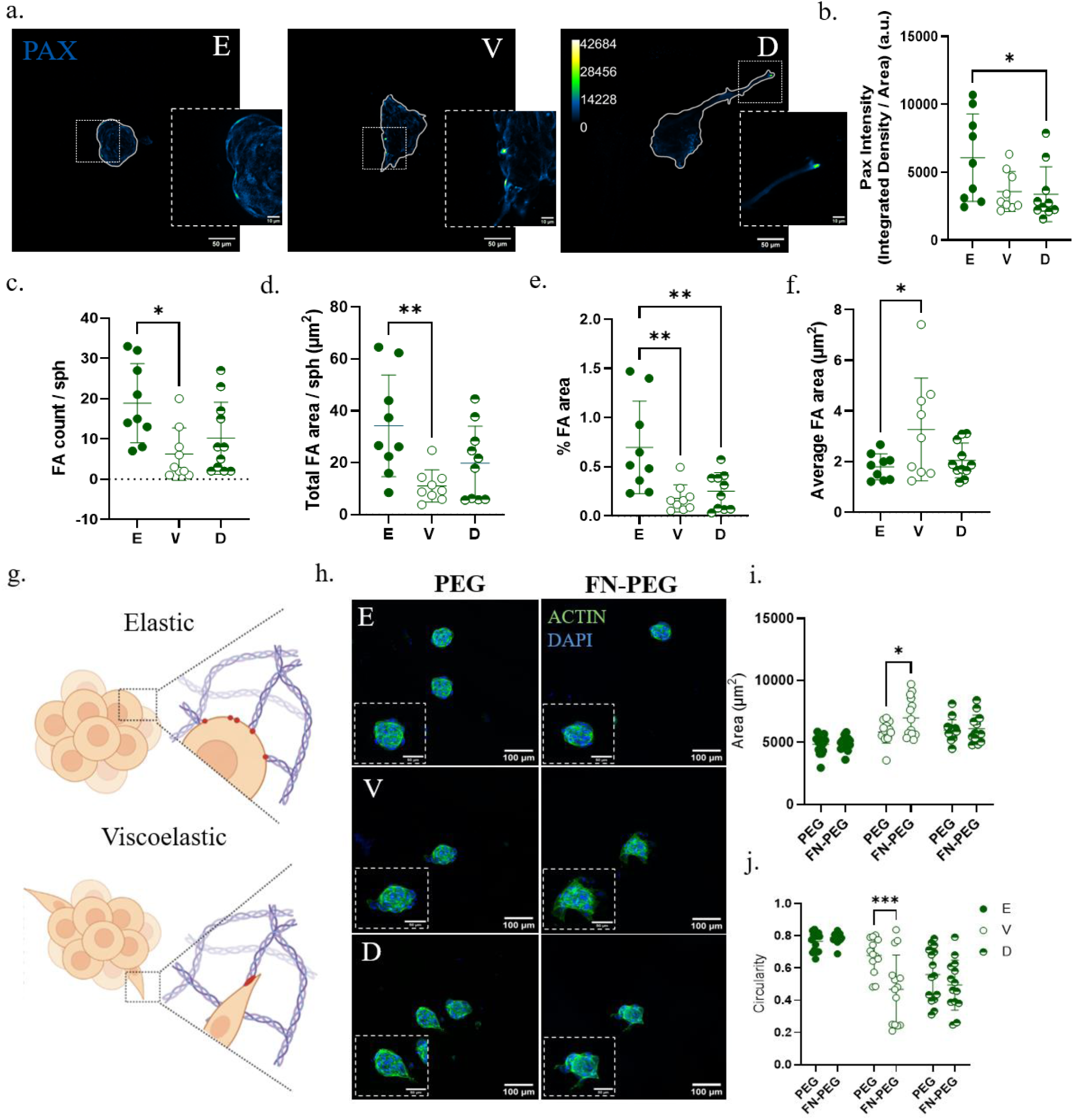
Spheroid adhesions are polarised and change depending on the mechanical environment. **(a)** Representative immunofluorescence images of hMSC spheroid paxillin-based adhesions in soft elastic, viscoelastic, and degradable hydrogels after 3 days. Scale bar = 50 μm in main panels and 10 μm in insets **(b)** Quantified paxillin intensity **(c)** focal adhesion (FA) count **(d)** total FA area **(e)** % FA area, and **(f)** average FA area in soft elastic, viscoelastic and degradable hydrogels. Data are shown as mean ± SD, SE: n = 9, SV: n = 9, SD: n = 10 spheroids per condition, N = 3. One-way ANOVA with Tukey’s post-hoc test for multiple comparisons. *\*p-value* < 0.05, *\*\*p-value* < 0.01 **(g)** Graphical representation of adhesion patterns in soft elastic vs viscoelastic hydrogels. Made in BioRender. **(h)** Representative immunofluorescence images of spheroids (actin cytoskeleton = green, DAPI = blue) encapsulated in soft PEG and FN-PEG elastic, viscoelastic, and degradable hydrogels after 24 hours. Scale bar = 100 μm in main panels and 50 μm in insets **(i)** Quantified spheroid area and **(j)** circularity in soft PEG and FN-PEG elastic, viscoelastic, and degradable hydrogels. Data is shown as mean ± SD, SE: n = 15, SV: n = 13-14, SD: n = 14-15 spheroids per condition, N = 2. Two-way ANOVA with Tukey’s post-hoc test for multiple comparisons. *\*p-value* < 0.05, *\*\*p-value* < 0.01, *\*\*\*p-value* < 0.001, *\*\*\*\*p-value* < 0.0001.

To determine whether spheroid spreading depend on FN-mediated cell-matrix interactions, spheroids were encapsulated in hydrogels with and without FN (FN-PEG and PEG, respectively) (**Fig. 3h**). Regardless of the presence or absence of FN, spheroid morphology remained unaffected in SE hydrogels, as evidenced by the absence of significant differences in spheroid area or circularity between PEG and FN-PEG hydrogels (**Fig. 3i,j**).

In SV hydrogels, by contrast, spheroid morphology was strongly influenced by the presence of FN. Spheroids in viscoelastic FN-PEG hydrogels displayed significantly larger projected areas and reduced circularity compared to those in viscoelastic PEG-only hydrogels. These morphological changes arose largely from the formation of multiple, tensioned protrusions, a morphological feature that was absent in PEG-only hydrogels, where spheroids remained more rounded and compact. (**Fig. 3i,j**). Notably, even without FN, spheroid area remained significantly greater in viscoelastic compared to elastic hydrogels, suggesting that the time-dependent mechanical properties of viscoelastic matrices alone are sufficient to promote a degree of collective spreading independent of FN-mediated adhesion (**Table S4**).

In SD hydrogels, however, spheroids exhibited spreading regardless of FN incorporation. Spheroid area and circularity did not differ between PEG and FN-PEG hydrogels, though both remained significantly different from those in the respective elastic counterparts (**Fig. 3i-j; Table S4**). These findings suggest that spheroids in degradable hydrogels can remodel and invade the surrounding matrix independent of exogenous FN in the hydrogel, potentially through interactions with their own secreted ECM. Supporting this, spheroids in degradable FN-PEG hydrogels produced significantly more cellular FN compared to spheroids in elastic hydrogels, and this FN appeared highly organised and fibrillar (**Supplementary Fig. 7a-d**), a result consistent with previous studies on single hMSCs encapsulated in proteolytically degradable norbornene-modified hyaluronic acid hydrogels^16^.

Together, these findings suggest that the requirement for FN-mediated adhesion in driving spheroid spreading and symmetry breaking is strongly dependent on matrix mechanical behaviour. Specifically, engagement with matrix-bound FN appears necessary to promote protrusive multicellular spreading in viscoelastic matrices, whereas matrix degradability alone is sufficient to support spheroid spreading independently of FN functionalisation.

### Matrix stiffness and degradability set the magnitude of collective traction forces

Traction forces are a primary mechanism through which cells sense and respond to the mechanical properties of the surrounding ECM. These forces are generated by the actomyosin cytoskeleton and transmitted to the ECM via focal adhesions or to adjacent cells via cadherin-based adherens junctions, a process which regulates a variety of cellular processes including migration and differentiation ^37^. Given the distinct morphological and focal adhesion patterns observed across different mechanical environments, we hypothesised that these structural differences would be accompanied by corresponding changes in the contractile forces generated at the spheroid-ECM interface. To test this, we incorporated fluorescent beads into soft (*G’* ≈ 0.5 kPa) and rigid (*G’* ≈ 1.5 kPa) elastic and degradable hydrogels of an established transglutaminase-crosslinked PEG platform to serve as fiducial markers for quantifying spheroid-induced hydrogel deformations and forces using 3D traction force microscopy (TFM; **Supplementary Fig. 8a-g**). The homogeneous network structure of this platform facilitated a uniform distribution of fluorescent beads throughout the matrix (**Fig. 4a**) and recapitulated the morphological behaviours previously observed (**Supplementary Fig. 9a-g**).Viscoelastic hydrogels were excluded from these experiments because their time-dependent stress relaxation complicates the reconstruction of bead displacement fields and precludes accurate traction force calculations using conventional 3D TFM algorithms, which assume linear elasticity.

**Figure 4:**
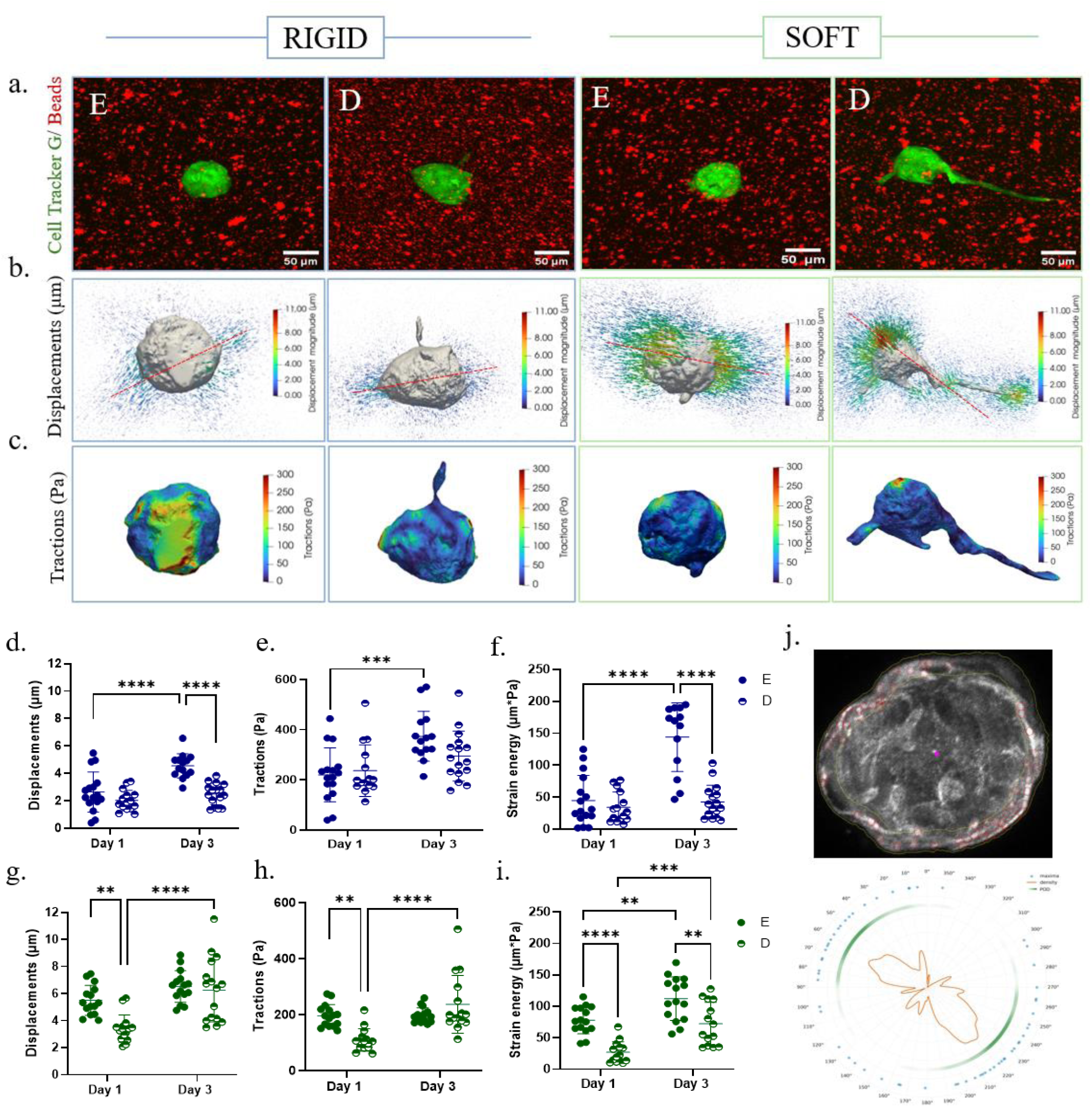
Spheroid-generated displacements and traction forces change depending on matrix mechanics and exhibit polarisation. **(a)** Representative maximally projected images of beads (in red) overlaid with green-labelled spheroids (top), **(b)** the corresponding 3D renders of displacements (middle) **(c)** and traction fields (bottom) computed around the spheroids after 3 days **(d-f)** Quantification of 99th percentile bead displacements, tractions and strain energy of spheroids encapsulated in rigid and **(g-i)** soft elastic and degradable hydrogels. Data are presented as mean ± SD, n = 16 spheroids per condition, N = 2. *\*p-value* < 0.05, *\*\*p-value* < 0.01, *\*\*\*p-value* < 0.001, *\*\*\*\*p-value* < 0.0001. Two-way ANOVA with Tukey’s post-hoc test for multiple comparisons. **(j)** Immunofluorescence image depicting paxillin maxima (red circels) and corresponding polar plot showing the angular distribution (0-360°) of paxillin maxima (blue dots), density of maxima (orange line) and product of opposite density (POD; green line).

As expected, both matrix mechanics and time significantly influenced MSC spheroid contractility. Matrix displacements and traction forces consistently oriented inwardly (arrows pointing towards the spheroid centre) across all conditions, with overall positive contractility values, suggesting that spheroids increasingly pulled rather than pushed on the matrix over time (**Supplementary Fig. 10a,b**). Spheroids in rigid hydrogels exhibited significantly smaller displacements but significantly higher forces compared to those in softer hydrogels at both time points, regardless of the degradation profile of the hydrogel (**Table S5)**. These results are consistent with previous observations in single fibroblasts encapsulated in synthetic PEG-MAL hydrogels, where increasing hydrogel stiffness resulted in lower matrix displacements but larger forces^38^. This indicates that MSC spheroids actively adapt their contractile behaviour to the stiffness of the surrounding matrix, generating stronger pulling forces in stiffer environments possibly to overcome the greater mechanical resistance of the matrix.

We next examined whether matrix degradability altered spheroid-generated forces within each stiffness regime. Spheroids in RE and RD hydrogels generated comparable displacements on day 1. However, by day 3 spheroids in RE hydrogels, which remained relatively rounded, generated over 50% higher matrix displacements relative to those in their degradable counterparts (**Fig. 4d**), reflecting a significant increase in displacements over time in RE hydrogels that was absent in RD hydrogels. Traction force and strain energy measurements revealed similar trends: spheroids in RE hydrogels exerted significantly higher forces on day 3 compared to day 1 (**Fig. 4e**), accompanied by a more than threefold greater strain energy than those in degradable hydrogels on day 3 and those in elastic hydrogels on day 1 (**Fig. 4f**). Thus, spheroids in rigid, elastic non-degradable matrices progressively increased force transmission to the surrounding ECM over time, despite no morphological change, a trend that was not observed for spheroids in stiffness-matched degradable hydrogels. These results highlight that spheroid morphology may not necessarily predict the magnitude of collective contractile activity. Furthermore, the reduced force generation and strain energy in degradable matrices may reflect the ability of cells to locally remodel and relieve mechanical stresses within the surrounding network, thereby reducing the accumulation of matrix tension over time. However, this interpretation remains speculative and would require direct measurements of matrix remodelling or local stress relaxation to confirm.

In softer hydrogels, the force dynamics were more complex. On day 1, spheroids encapsulated in elastic hydrogels generated significantly larger matrix displacements and tractions than those in degradable hydrogels (**Fig. 4g,h**) but this difference disappeared by day 3, as spheroids in degradable hydrogels exhibited a significant increase in both matrix displacements and tractions over time. Notably, this rise in force generation coincided with the pronounced spheroid spreading observed in SD hydrogels (**Fig. 4a**), suggesting that enhanced collective spreading is accompanied by increased contractile activity, a result consistent with previous studies in single cells^39,40^.

Analysis of the strain energies experienced by the hydrogels further supported the time-dependent effect of degradation (**Fig. 4i**). On day 1, the strain energy in elastic, non-degradable hydrogels was significantly greater than that in degradable hydrogels. Although strain energy increased significantly over time in both conditions, values remained significantly higher for elastic hydrogels at day 3 despite comparable matrix displacements and forces between the two groups. This likely reflects the greater mechanical resistance of the non-degradable matrix, which allows greater elastic energy to accumulate for a given level of cellular force generation.

Together, these findings reveal a stiffness-dependent and time-dependent effect of matrix degradation on collective force generation. In rigid matrices, degradability was associated with reduced accumulation of matrix displacement, traction force, and strain energy over time, despite minimal changes in spheroid morphology. In contrast, in soft matrices, degradability initially reduced force generation but was followed by an increase in displacement and traction force coinciding with pronounced spheroid spreading. Thus, degradability does not simply suppress collective contractile activity; rather, its effect depends on the mechanical context of the matrix and the timescale of the cellular response. These findings further demonstrate that morphological remodelling and mechanical output can become decoupled, with spheroids capable of altering their contractile interaction with the ECM even without substantial changes in overall morphology.

### The force field and cell-matrix adhesions are polarised along a single axis, independently of spheroid shape

Reconstruction of 3D bead displacement fields and traction force maps also revealed a consistent polarisation of high-magnitude matrix deformations and tractions, which typically localised to opposite sides of the spheroid surface (**Fig. 4b,c; red lines**). Critically, this pattern was present in spheroids in non-degradable elastic hydrogels that remained spherical as well as in those that had broken symmetry. The force field is therefore polarised before, and independently of, any change in spheroid shape. These observations suggest, first, a directional organisation of collective force generation that persists across spheroid morphologies and matrix conditions and, second, that once symmetry is broken, contractile activity is redistributed so that force generation increases at the pole opposite the emerging protrusion, a behaviour previously reported for single cells in 3D environments^39^. Such spatial reorganisation could contribute to subsequent protrusion formation and thereby facilitate redistribution and rebalancing of forces across the spheroid.

We next asked whether cell-matrix adhesions share this spatial organisation around the spheroid by analysing the distribution of paxillin along the spheroid periphery. Because the angular analysis requires a well-defined, approximately circular boundary, only spheroids in elastic hydrogels analysed. Paxillin maxima within a narrow rim region, specifically selected to assess cells in direct contact with the surrounding hydrogel, were identified and mapped in angular coordinates (**Fig. 4j**). In most spheroids, paxillin maxima were not uniformly distributed around the spheroid rim, but instead concentrated within two broad, approximately opposing angular regions. This organisation was further supported by density profiles and by the product-of-opposite densities (POD) metric, which indicated a dominant axis of paxillin organisation around the spheroid perimeter. Collectively, these findings indicate that multicellular aggregates preferentially organise contractile forces and cell–matrix adhesions along discrete opposing poles **(Supplementary Fig. 11)**, consistent with the polarised force patterns observed in 3D TFM.

### An anisotropic distribution of contractile cells is sufficient to generate the polarised force field

To identify the mechanism underlying the polarised displacements observed experimentally, we developed a simplified computational model describing a spheroid embedded within an elastic, non-degradable block of hydrogel matrix. The aim was to test whether internal cellular heterogeneity, in the form of an increased density of contractile cells along one axis, could generate anisotropic matrix deformations resembling those measured by 3D TFM. A schematic of this central hypothesis is shown in **Fig. 5a**, which contrasts the expected displacement patterns for spheroids with homogeneous versus heterogeneous cellular distributions. A uniform distribution of contractile cells within the spheroid is expected to produce isotropic, radially symmetric deformations, whereas a higher cell density along one axis generates anisotropic, polarised displacements aligned with that axis.

**Figure 5:**
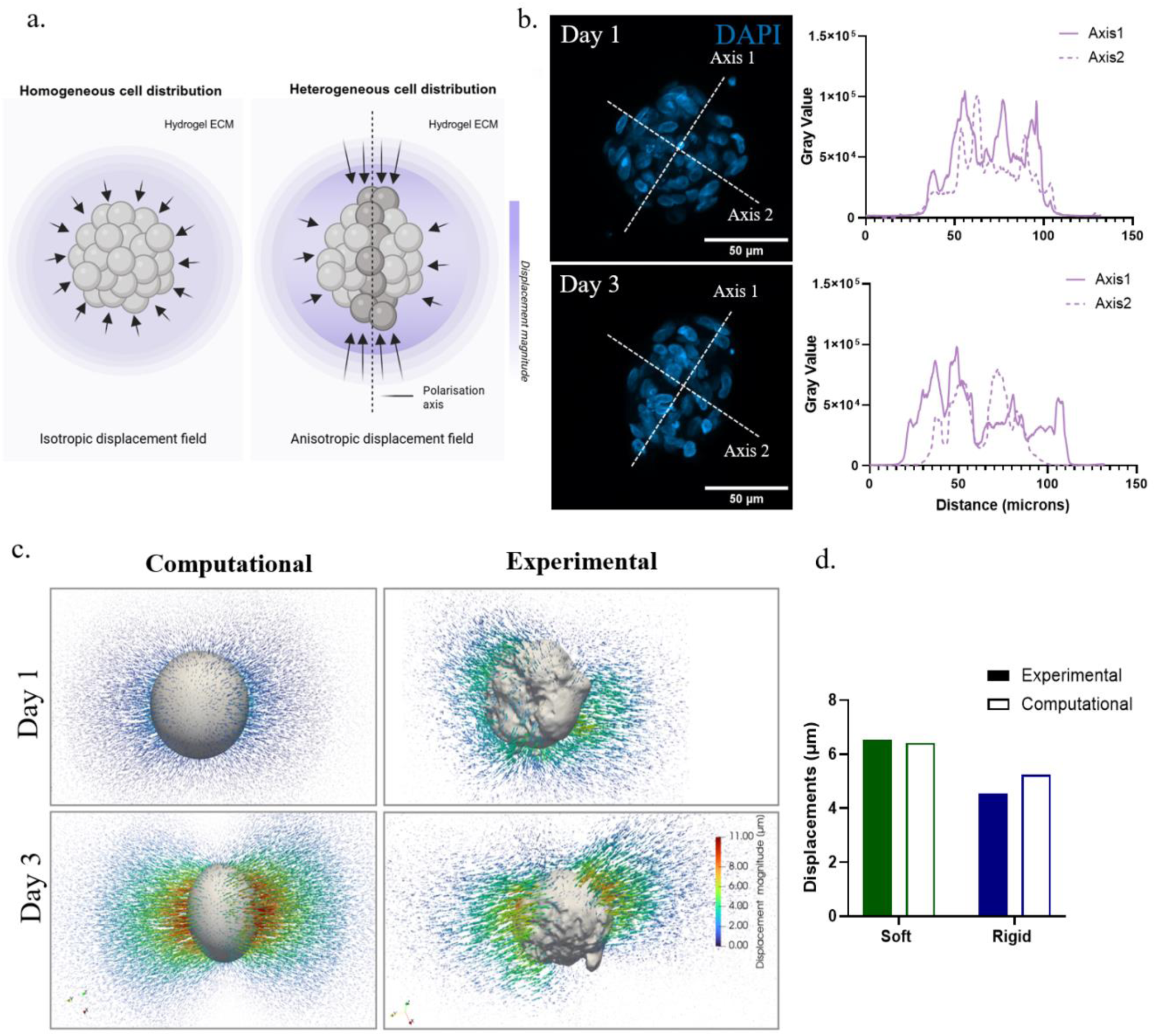
Experimental and computational spheroid-generated matrix displacement fields for spheroids in soft elastic hydrogels. **(a)** Schematic illustrating the core concept of the computational model. In the first panel (left), a spheroid with a homogeneous cell distribution generates uniform, radially symmetric displacements in the surrounding hydrogel. In the second panel (right) the spheroid contains a higher density of cells along one axis (heterogeneous distribution) producing anisotropic, polarized contractility. **(b)** Representative DAPI z-projections of spheroids encapsulated in non-degradable soft elastic hydrogels on day 1 and day 3 along with corresponding DAPI intensity profiles along two perpendicular axes (∼130 μm length) passing through the spheroid center. Scale bar = 50 μm **(c)** Visual comparison between the experimental (left) and computational displacement fields (right) for spheroids embedded in soft elastic hydrogels at day 1 and 3. **(d)** Quantitative comparison of the average 99th percentile displacements (μm) between experimental data and computational model for soft and stiff hydrogels on day 3.

This hypothesis was motivated by experimental observations of nuclear distributions within spheroids **(Fig. 5b)**. On day 1 spheroids exhibited comparable fluorescent profile widths along both axes tested, consistent with an isotropic nuclear distribution. In contrast, on day 3, spheroids displayed a broader nuclear distribution along axis 1 compared to axis 2, suggesting an anisotropic, preferential organisation of nuclei along axis 1. These observations informed both the formulation of the model and the choice of anisotropic growth parameters used in the simulations.

The computational model reproduced the key spatial features of the experimentally measured displacement fields. At day 3, both the experimental and computational results exhibited a pronounced orientation of matrix displacements along a principal direction, rather than being uniformly distributed around the spheroid **(Fig 5c)**. Importantly, the anisotropic displacement pattern emerged only when an uneven cellular distribution was imposed along a single axis, indicating that directional differences in cell density are sufficient to generate polarised matrix deformations similar to those observed experimentally. In contrast, the displacement field on day 1 appeared considerably more homogeneous. At this earlier stage, the cells presumably have not yet developed a pronounced preferential orientation, and their contractile activity is therefore distributed more uniformly throughout the spheroid. Consequently, the deformation induced in the surrounding hydrogel is more isotropic **(Fig. 5c)**. Therefore, the transition from a relatively homogeneous displacement field on day 1 to a preferentially oriented deformation pattern on day 3 is consistent with the progressive reorganisation and alignment of the cells within the spheroid over time.

In addition to capturing the spatial organisation of the displacement fields, the model also reproduced their magnitudes across both hydrogel stiffnesses. The upper range of the displacement distributions (99^th^ percentile) showed close agreement between simulations and experiments for soft hydrogels, while displacements were slightly overestimated for stiff hydrogels **(Fig. 5d).** Additional simulations confirmed that displacement magnitudes were largely insensitive to the Poisson’s ratio of the hydrogels, and that comparable magnitudes could be obtained through different combinations of spheroid stiffness and contractility **(Supplementary Fig. 12-13),** indicating that the model does not rely on a narrow parameter regime to reproduce the observed behaviour.

Together, these results show that this simplified anisotropic-contractility model can recapitulate both the spatial organisation and magnitude of spheroid-induced matrix deformations. This suggests that internal cellular heterogeneities within the spheroids, arising during early collective reorganisation or symmetry breaking, are sufficient to drive the emergence of polarised force generation in multicellular aggregates. This provides a candidate mechanism linking internal organisation to subsequent protrusion formation and directional migration.

## Discussion

The mechanical properties of the ECM, including stiffness, viscoelasticity, and degradability are increasingly recognised as key regulators of collective cell behaviour. Yet these properties are usually either studied separately or coupled within a single material system, making their distinct roles difficult to resolve. Here, by independently tuning these properties in 3D PEG-based hydrogels, we find that multicellular spheroids intrinsically develop mechanical polarity, while matrix properties govern whether this polarity is translated into morphological asymmetry. The spheroid carries a mechanical axis, visible in its traction field and adhesion organisation, even when its shape remains spherical. However, the mechanics of the surrounding network set whether this axis is converted into a change in morphology.

Stress relaxation and degradability were not equivalent: viscoelasticity promoted symmetry breaking in soft hydrogels but not in rigid ones. This behaviour is consistent with the idea that stress relaxation can reduce the mechanical resistance encountered by cells and thereby permit morphological change when the surrounding network is sufficiently compliant. Previous studies have shown that stress relaxation can promote spreading of individual cells and multicellular aggregates in compliant matrices, results that agree with the data shown here^27,29,34,41,42^. However, the magnitude and kinetics of this response as well as the initial stiffness of the environment vary substantially between hydrogel systems, reflecting differences in network architecture, ligand presentation and, importantly, the timescale and extent of stress relaxation.

Degradability produced a more pronounced response. Proteolytic cleavability of the network enabled spreading regardless of matrix stiffness, indicating that local loss of network connectivity is sufficient to permit spheroid symmetry breaking and protrusion formation. Nevertheless, the extent of spreading remained sensitive to matrix stiffness, with stiffer degradable matrices imposing greater resistance to spheroid outgrowth. This distinction is consistent with previous work on MSCs and endothelial cells suggesting that matrix stiffness can influence the final extent of cell spreading even when similar levels of degradability are present^43,44^. At the collective level, our findings extend this concept by showing that stress relaxation and degradation can both facilitate morphological change, but through fundamentally different physical mechanisms: stress relaxation dissipates stored mechanical energy, whereas degradation locally removes the network that constrains cellular rearrangement.

This distinction may be particularly important for multicellular systems. A spheroid cannot simply extend a single cell membrane into a deformable matrix; expansion requires coordinated changes in cell–cell contacts, cell–matrix adhesions and the transmission of forces across many cells. Thus, the requirement for matrix compliance in viscoelastic conditions, but not degradable conditions, suggests that the ability of the matrix to accommodate collective deformation is a key determinant of whether internal cellular forces translate into changes in the macroscopic spheroid morphology.

Furthermore, these matrix-dependent morphological changes were accompanied by changes in spheroid core stiffness. Brillouin measurements revealed lower frequency shifts in more spread spheroids encapsulated in viscoelastic and degradable hydrogels, compared with the higher frequency shifts in circular spheroids embedded in elastic hydrogels. Similar matrix-dependent Brillouin responses have been reported in MCF-7 cancer spheroids, with those encapsulated in degradable PEG-heparin hydrogels exhibiting lower frequency shifts compared to those in stiffness-matched non-degradable hydrogels^36^. One possible explanation is that matrix-permissive conditions alter morphology, leading to different levels of cellular packing and increased intercellular spaces during spreading, thereby modifying the spheroid-level mechanical response. This suggests that multicellular aggregates act as active entities whose mechanical state is not fixed but can instead adapt to the mechanical properties of the surrounding matrix through changes in cell rearrangement, cytoskeletal remodelling and changes in cell–cell organisation.

These observations connect two measurements that are otherwise reported separately. Brillouin microscopy probes the mechanical field inside the aggregate, whereas traction force microscopy probes the field it imposes on the matrix, and both lose radial symmetry under the conditions that permit symmetry breaking. The computational model offers a direct link between them: if an anisotropic distribution of contractile cells generates the polarised displacement field, the same anisotropy should be visible as an asymmetry in the internal mechanical profile, since the Brillouin frequency shift is sensitive to cell packing. This predicts that the axis of Brillouin asymmetry and the axis of traction polarity should coincide within a given spheroid.

Cell-matrix adhesions provide a potential cellular mechanism underlying these collective responses. Although integrin-mediated cell-matrix adhesions are well established in 3D^33,45^, evidence for the formation of canonical focal adhesions comparable to those observed on 2D substrates remains mixed. Here, paxillin-positive adhesion structures were detected across all matrix conditions, regardless of morphological characteristics, but their organisation and size changed substantially with matrix mechanics. Spheroids in elastic hydrogels were associated with smaller adhesions that occupied a greater cumulative adhesion area, consistent with a more extensive peripheral cell-matrix adhesion. In contrast, viscoelastic hydrogels promoted fewer, larger adhesions, particularly around protrusive regions. These patterns agree with reports that viscoelasticity promotes the formation of intense, punctate FA clusters, and are consistent with the established sensitivity of adhesion assembly and maturation to mechanical loading^33^. However, whether this represents altered adhesion turnover, maturation or force transmission remains to be established.

A related principle may apply to degradable matrices. Local network cleavage provides cells with an additional mechanism for relieving mechanical constraints around adhesions and extending protrusions. Consequently, productive collective remodelling may not require extensive stabilisation of adhesions across the entire spheroid boundary. Instead, force transmission may become concentrated at a smaller number of mechanically active regions where matrix engagement and remodelling occur simultaneously.

The 3D traction force measurements further demonstrated that matrix mechanics regulate the mechanical output of the spheroid. We found that traction forces increased with matrix stiffness, consistent with established mechanotransduction mechanisms in single cells in 2D and 3D environments^39,46,47^. However, degradability produced a more context-dependent response whereby both the mechanical environment and the stage of spheroid adaptation played an important role.

This behaviour is consistent with the idea that matrix degradation changes the mechanical boundary conditions experienced by the spheroid. By locally removing load-bearing material, degradation can reduce the requirement for high-magnitude forces to generate deformation, while simultaneously creating new regions into which cells can extend and remodel. Thus, the resulting force magnitude reflects a balance between cellular contractility and the evolving resistance of the surrounding matrix, rather than a direct response to matrix stiffness alone.

The time dependence of force generation can be read in the same way. In rigid elastic hydrogels, spheroids increased matrix displacements, traction force and strain energy between day 1 and day 3 without any change in shape. If the force field is already polarised in these spheroids, this growing mechanical output would be transmitted through the same two poles rather than being distributed over the aggregate surface, and the polarity should therefore strengthen with time.

Perhaps most strikingly, tractions were not distributed uniformly around the spheroid but were spatially polarised. Regions of elevated tractions and matrix deformations emerged at approximately opposing locations, accompanied by a related polarisation in focal adhesion organisation. Although the adhesion measurements were obtained from two-dimensional projections, in a different hydrogel platform from the traction measurements, and therefore cannot fully resolve the three-dimensional organisation of adhesions, their spatial correspondence with the traction pattern suggests that force transmission is organised at the collective scale. A simplified computational model showed that anisotropic distribution of cells within the spheroid alone is sufficient to generate polarised displacement fields resembling those observed experimentally. Previous observations have shown that multicellular aggregates can undergo collective transitions from orbiting to matrix invasion, demonstrating that spatial organisation within a multicellular aggregate can strongly influence how forces are transmitted to the ECM^26^. Our results extend this concept by linking internal organisation to the *spatial* pattern of forces transmitted to the surrounding ECM. The spheroid therefore acts not simply as a collection of mechanically active cells, but as a mechanically coupled entity in which local heterogeneities in cell organisation can generate patterns of force transmission and possibly symmetry breaking. The origin of this anisotropy remains, however, open: it could arise stochastically during spheroid formation and be amplified by mechanical or biochemical feedback or emerge from active cell alignment during the initial day of culture. These possibilities are not mutually exclusive, and distinguishing them will require further research, possibly by tracking cell positions from spheroid formation onwards.

Our results support a picture in which force polarity is an intrinsic property of multicellular aggregates and matrix mechanics act as a filter on its expression. More broadly, they establish stiffness, viscoelasticity and degradability as distinct mechanical regulators of collective mechanobiology that can differentially regulate spheroid responses such as morphology, stiffness, adhesion organisation and force generation. Notably, our work shows that at the multicellular level, cell-cell interactions, spatial organisation and collective force transmission introduce additional degrees of freedom through which mechanical information is integrated. Understanding how these responses emerge or are coupled may be important in physiological and pathological settings in which cells encounter matrices that simultaneously exhibit many of these properties, including in processes such as development, tissue repair and tumour progression. More generally, independent control of these properties in engineered matrices offers a route to directing collective cell behaviour and should aid the development of more physiologically relevant in vitro tissue models and regenerative therapies.

## Materials and Methods

### Cell culture

Cryopreserved human bone marrow-derived mesenchymal cells (hBM-MSCs) (Promocell) were maintained in a 1:1 ratio of Dulbecco’s Modified Eagle’s medium (DMEM) (Sigma) (containing 1% Sodium Pyruvate (Sigma), 1% MEM NEAA (non-essential amino acids, Gibco), 2% of a mixture of L-glutamine (Gibco), Penicillin/Streptomycin (Sigma), and Amphotericin B (Gibco)) and Mesenchymal Stem Cell Growth Medium 2 (with supplement mix (Promocell)). Cells were incubated at 37°C in a humidified environment (5% CO_2_). After a day, the culture medium was changed to DMEM supplemented with 10% Fetal Bovine Serum (FBS; Gibco). The medium was changed thereafter every two days until cells reached ∼80% confluence, at which point the cells were trypsinised with pre-warmed 0.05% trypsin-EDTA (v/v) and counted with a haemocytometer in the presence of trypan blue (Gibco).

### Spheroid culture

Spheroids were generated using AggreWell™400 24-well plates (StemCell Technologies) pre-treated with anti-adherence rinsing solution according to the manufacturer’s instructions. Wells were centrifuged (1300 ×g, 5 min), washed twice with PBS, and filled with 200 μl DMEM without FBS prior to incubation. Trypsinised cells (120,000 cells per well, yielding ∼100 cells per spheroid) were resuspended in DMEM without FBS to a final volume of 500 μl per well, centrifuged (100 ×g, 3 min), and incubated for 24 h. Spheroids were then washed with PBS and collected by gentle resuspension. All collection plastics were pre-coated with 2% FBS in PBS. Spheroids were pelleted (5 min, room temperature), resuspended in 1 ml PBS, and quantified by manual counting from a 50 μl aliquot.

### Fibronectin PEGylation

Fibronectin (FN) was PEGylated according to a previously published procedure^28^. First, FN (YoProteins; 3 mg/ml) was denatured in PBS containing 5mM Tris(2-carboxyethyl)phosphine hydrochloride (TCEP, pH 7, Thermo Scientific) and 8 M Urea (Sigma, U5128, 99.5%) for 15 min at RT (20-22°C). 4-arm-PEG-Maleimide (PEG-Mal, 20 kDa, Creative PEGWorks) was then added at a molar ratio of 1:4 FN:PEG-Mal. The reaction was quenched with 2 μl NaOH (1M stock) and unreacted cysteine residues were blocked with 14 mM iodoacetamide (Sigma) (pH 8 for 2 hours). Excess reagents were removed by dialysis against PBS for 1h at RT, and the protein solution was precipitated in cold absolute ethanol overnight at -20°C. The solution was then pelleted by centrifugation at 7,150×g / 4°C for 30 min, washed with 90% cold ethanol and centrifuged again for 20 min. After removing the ethanol, pellets were left to fully dry, and FN was resuspended in 8 M urea to 3 mg/ml. Once dissolved, the solution was dialysed for 1h at RT against PBS and stored at -20°C until use.

### 3D hydrogel preparation

#### PEG-MAL hydrogels

PEG–Mal hydrogels (50 μl) were formed via Michael-type addition reaction at physiological pH (7.4). PEG–Mal and thiolated crosslinkers were dissolved in PBS (pH 6–6.5), and PEGylated fibronectin (FN–PEG) was added to the PEG–Mal solution at a final concentration of 0.5 mg ml⁻¹. Crosslinking was initiated by mixing the PEG-Mal/FN-PEG solution with thiolated crosslinkers at a 1:1 maleimide:thiol molar ratio. The crosslinkers used were either PEG-dithiol (non-degradable crosslinker) or a 1:1 mixture of PEG-dithiol and an MMP-sensitive peptide crosslinker (VPM). Cells or spheroids were included in the PBS component before mixing. Gels were cast between hydrophobic surfaces, with the top one being a treated coverslip (RainX-treated, 9 mm), and allowed to polymerise for 30 min at 37 °C before transferring to 24-well plates and incubation in DMEM with 10% FBS.

#### PEG-VS hydrogels

PEG–VS degradable and non-degradable hydrogels (2 & 3% w/v for soft and rigid, respectively; 50 μl) for 3D traction force microscopy were prepared as previously described^48^. Briefly, PEG hydrogels were enzymatically crosslinked using coagulation enzyme-activated transglutaminase factor XIIIa (FXIIIa) between a glutamine peptide-modified PEG precursor (8arm, 40kDa) and either a non-degradable lysine peptide-modified PEG precursor (AcFKGG-GDQGIAGF-ERCG-NH2 (peptide A)) or an MMP-sensitive lysine peptide-modified PEG precursor (AcFKGG-VPMSMRGG-ERCG-NH2 (peptide V)). Far-red fluorescent beads (200 nm) at a 1:60 concentration, MilliQ water, 10X buffer, PEG precursors, lysine-RGD (50 μM), and spheroids in PBS, were first mixed in this order, followed by FXIIIa, which activated crosslinking. Spheroids were previously stained with Cell Tracker Green (ThermoFisher) at a 1:1000 concentration for 30 minutes at 37°C. After the addition of FXIIIa, pre-polymerised solutions were pipetted into a 24-well glass-bottomed well-plate (Cellvis) and left to crosslink at 37°C for 30 minutes, after which they were submerged in DMEM with 10% FBS.

### Rheology

The bulk mechanical properties of PEG-Mal hydrogels (100 μl) were measured using a stress-controlled rheometer (MCR302, Anton Paar) with a parallel plate geometry (upper plate diameter 8 mm) at 25°C. Samples were hydrated with PBS prior and during testing. Strain sweep tests were performed at a constant angular frequency of 10 rad/s, over a strain range of 0.1% to 10% strain, to identify the linear viscoelastic regime (LVE) of the gels. Frequency sweep tests were conducted at a range of 1 to 100 rad/s (0.1 – 10 Hz) at a strain of 1% or 10%, depending on the LVE of the hydrogels. Storage and loss moduli, *G’* and *G’’* were determined at a gap size corresponding to a normal force of ∼0.1 N. Stress relaxation was assessed by investigating the decrease shear stress over 932 s at a constant strain of 1% or 10% (depending on the LVE profile observed during strain sweeps).

Stress relaxation data were analysed using a custom analysis script in the form of a Jupyter notebook (Code publicly available on GitHub; https://github.com/GiuseppeCiccone96/stressrelaxbulkrheo). Shear stress (σ) data from the first second of measurements were excluded to minimise noise. And stress values from 1-932 s were averaged across technical replicates and normalised to the initial stress (σ/σ0) to obtain an average stress relaxation curve per condition. Average stress relaxation curves were used to quantify the time required to relax 15% of the initial stress; where this was not achieved, the final time point (932 s) was reported.

### Brillouin Microscopy

Brillouin microscopy experiments were conducted as previously described elsewhere^49^. Briefly, at days 1 and 3, samples were transferred to glass-bottom dishes prior to Brillouin spectroscopy measurements. Measurements were performed using an inverted light Brillouin microscope (LightMachinery) equipped with a two-stage VIPA-based spectrometer (Hyperfine, LightMachinery). Samples were illuminated with a 660 nm single-mode laser at ∼50 mW using a 20× (NA 0.45) objective at room temperature. Backscattered light was collected through the same objective, spectrally dispersed by the VIPA spectrometer, and detected using a high-sensitivity camera (Orca Fusion).

Spectrometer calibration and background subtraction were performed prior to acquisition using an acrylic cube. Brillouin maps were acquired using stage-scanning controlled by custom LabVIEW software (LightMachinery), generating cross-sectional maps through the spheroid centre in the XY plane. Individual spectra were fitted with Lorentzian functions, and Brillouin frequency shifts, V_B_, were extracted relative to the Rayleigh peak. Maps covered 120–180 μm (spheroids) or 100 μm (empty hydrogels) in X and Y with a 7 μm step size.

Finally, Brillouin spectra were analysed using a custom Python script (Code publicly available on GitHub; GiuseppeCiccone96/brillouinmaps) in combination with ImageJ (v2.14.0). Colour-coded Brillouin frequency shift maps were first generated with appropriate x and y coordinates. Brightfield images were then used to manually segment regions of interest (ROI; e.g. spheroid area). Segmentation masks were resized to match Brillouin map resolution and binarised prior to analysis. Brillouin frequency shifts within the segmented ROIs were extracted to generate spatial profiles and colour maps, and the mean Brillouin frequency shift ± standard deviation was reported for each ROI.

### Immunofluorescence

Samples were fixed with 4% paraformaldehyde in PBS for 30 minutes at RT. Following fixation, samples were washed twice with PBS and permeabilised with 0.1% (v/v) Triton X-100 (Sigma) in PBS for 30 minutes. Samples were blocked in 1% (w/v) Bovine Serum Albumin (BSA; Sigma) in PBS for at least 1h at RT and then incubated overnight at 4°C with the relevant primary antibody. Primary antibodies used were Anti-human Fibronectin (ThermoFisher, 1:150) and Ki-67 (Cell Signaling, 1:250). Samples were subsequently washed twice for 15 minutes with 0.1% (v/v) Tween-20 (Sigma) in PBS and once with PBS for 10 minutes on the shaker. Secondary antibodies and phalloidin (1:250), which stains the actin cytoskeleton, were then added for 1h and left shaking at RT in the dark. Hydrogels were then washed twice for 5 minutes with 0.1% Tween-20 in PBS and once for 10 minutes with PBS while shaking. An appropriate volume of PBS containing DAPI (1:400 dilution) was added to each sample for 20 minutes to stain nuclei. Gels were then washed with PBS twice for 5 minutes and left in PBS at 4°C until imaged.

Confocal fluorescent microscopy was carried out using an LSM-980 confocal microscope (Zeiss). Confocal images were acquired using a 10x 0.3 NA or a 20x 0.8 NA dry objective at a 1982 x 1982 resolution, using an excitation/ emission wavelength of 493 nm/ 517 nm for the green channel which was typically used for visualisation of the actin cytoskeleton, excitation/emission of 548 nm/ 561 nm for visualisation of FN staining, excitation/emission of 647 nm/ 668 nm for visualisation of paxillin staining, and excitation/emission of 353 nm/ 465 nm respectively for DAPI nuclear staining. Within experiments and amongst replicates, imaging settings were kept constant. Z-stacks spanning the entire spheroid were acquired with a step size of ∼5-10 μm.

### Image analysis

#### FN functionalisation analysis

Fluorescent FN hydrogel images were analysed using ImageJ2 (version 2.14.0; National Institute of Health, USA). Each hydrogel condition was prepared in duplicate, and for each hydrogel, two images were acquired. Z-stacks were collapsed using maximum intensity projection, and a square region of interest (ROI) was used to measure the integrated density in three areas per image. To account for background signal, a negative control (FN-PEG hydrogel without Alexa-488 labelling) was processed identically. Reported FN intensity values were calculated by averaging the integrated density of the three measured areas per image, then subtracting the average integrated density of the negative control.

#### Spheroid morphology analysis

Cell and spheroid shape descriptors, such as area, circularity and aspect ratio, were measured using ImageJ 2.14.0v (National Institute of Health, US). Briefly, z-stacks were collapsed using maximal intensity projection and channels were split. A Gaussian Blur of sigma 2 was applied to the phalloidin channel (staining the actin cytoskeleton) prior to thresholding each cell or spheroid. Using the wand tracing tool, thresholded features were selected and the resulting ROIs were used to quantify shape descriptors of the enclosed areas.

#### Cellular FN production analysis

Fibronectin production was analysed using ImageJ 2.14.0v (National Institute of Health, US). Cell-secreted FN was stained using an anti-human fibronectin monoclonal antibody (ThermoFisher) that recognises a determinant on human cellular but not plasma fibronectin. Once imaged, Z-stacks were collapsed using maximal intensity projection and channels were split. A Gaussian Blur with a sigma of 2 was applied to the phalloidin channel before thresholding each spheroid. Using the wand tracing tool, thresholded spheroids were selected, and the resulting ROIs were used to quantify FN intensity on the FN channel. To quantify and subtract background intensity, a ROI was created and used to measure background intensity of the FN channel at two different areas of each image in all experimental conditions. The average of the mean gray value of all background measurements was then used to calculate FN intensity for each hydrogel condition using the Corrected Total Cell Fluorescence (CTCF) formula:

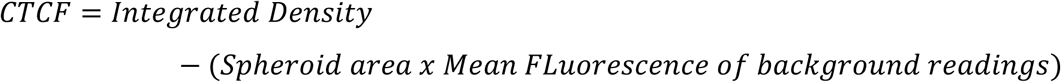

Finally, to identify any non-specific binding from the secondary antibody and validate signal intensity in experimental conditions, a secondary-only control sample was prepared for each hydrogel condition. These samples were only stained with the secondary antibody and imaged using the same imaging parameters as the experimental conditions.

#### Focal adhesion analysis

FA quantification was performed by adapting a protocol previously described in the literature^47^. Briefly, image scale was calibrated to pixel size, and paxillin channel images were converted to 8-bit. Background was subtracted (rolling ball radius = 50), and a Gaussian Blur was applied (sigma = 0.5). Contrast was enhanced using the CLAHE plugin (block size = 19, histogram bins = 256, maximum slope = 3, mask = none), and a mathematical exponential transform (Process > Math > Exp) was applied to further minimise background signal, followed by a second contrast enhancement step (0.35% saturation). A fixed threshold (70-255), determined by histogram observation, was applied uniformly across all images to generate a binary mask. Particles smaller than 0.5 μm² were excluded from the analysis. The results were saved both for individual focal adhesions as well as a summary of all particles in the spheroid.

#### Focal adhesion distribution analysis

The volumetric image data of the speroids were maximally projected into 2D images. A band of thickness ∼6 μm was defined around the circumference of each spheroid image. Intensity maxima in the band were obtained with a maximum spatial filter, using scikit-image’s <u>peak_local_max</u> function^50^. Finally, the angular coordinate of each maximum point was computed, using the centroid of the respective spheroid image as a pole.

The angular density of the maxima was computed with kernel density estimation, using a Gaussian kernel. To further analyse polarity, a new metric called product of opposing densities (POD) was defined. It was derived by multiplying the density estimates at two opposing angles, i.e. with 180° separation between them, and then normalising to the range [0, 1] across all angles.

#### Ki-67 analysis

The percentage of Ki-67-positive-cells for soft elastic, viscoelastic and degradable hydrogels was quantified using ImageJ 2.14.0v (National Institute of Health, US). Briefly, Z-stacks of spheroids stained with DAPI and Ki-67 were collapsed using maximal intensity projection and channels were split. The DAPI channel was used to manually count the number of total nuclei per spheroid while the Ki-67 channel was used to count the number of proliferating (Ki-67-positive) nuclei, respectively. The percentage of Ki-67-positive cells were then calculated using the formula:

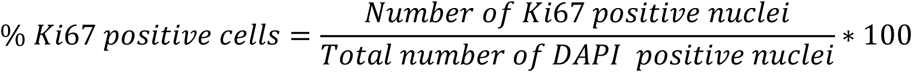

### 3D Traction Force Microscopy

Confocal images were acquired using a Leica SP8 Dive microscope equipped with a temperature-controlled stage maintained at 37°C and a humidified 5% CO2 environment. At 24 and 72 hours of culture, images were acquired at a 512 x 512 resolution using a 20x 0.4 NA dry objective. The pinhole was set at 1 AU with a diameter of 56.6 μm, and bi-directional imaging with phase correction was employed. The excitation wavelengths used were 488 nm (green channel for cells) and 638 nm (far-red channel for beads), which were detected with PMT detectors at 498-538 nm and 673-704 nm emission wavelengths, respectively. For each condition, all positions of interest were initially identified and marked. The selected positions were distributed across the hydrogel and spanned various depths along the z-axis. To capture displacements, images of spheroids were initially acquired when cells were mechanically active (stressed state), followed by treatment with Cytochalasin D (dissolved in DMSO, Sigma Aldrich) at a final concentration of 10 μM for 1 hour at 37°C, which mechanically inactivated the cells. Once contractility was disrupted (relaxed state), images of the same positions were acquired.

Spheroid 3D TFM data were analysed using the software toolbox TFMLAB (Publicly available at https://gitlab.kuleuven.be/MAtrix/Jorge/tfmlab_public)^51^. Briefly, the TFMLAB workflow performs the following operations: first, bead and spheroid images were filtered, and their contrast was enhanced. Spheroid images were further binarised to obtain a 3D mask. Second, using phase correlation, any microscope stage drifts that were present between the stressed and relaxed states were corrected. Third, free-from deformation-based image registration was used to measure the 3D bead displacements ^52^. ParaView (ParaView Software) was used to visualise bead displacements for each spheroid. Finally, tractions exerted by the spheroids were calculated from the measured displacements using a physically-based inverse method^53,54^. This method was developed in the framework of Finite Elements, and it computes a displacement field similar to the measured one while ensuring equilibrium of forces in the hydrogel domain. Due to the linear elastic nature of PEG-VS hydrogels used in these experiments, the hydrogel domain was modelled as a linear elastic material, with Poisson’s ratio ν = 0.3. The storage moduli, G’, were converted into Young’s moduli, E’, through the following equation for all conditions measured.

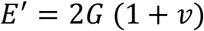

TFM results were postprocessed using in-house MATLAB code to quantify several metrics of interest. Specifically, the metrics quantified included:

1. 99th percentile traction (Pa) or displacement (µm): Given the traction field, ***t***, at the *N* nodes of the FE mesh, the values of the traction 2-norm,‖***t****n*‖2, are first sorted and then taken as the 100(0.5/N)th, 100(1.5/N)th, …, 100([N – 0.5]/N)th percentiles. For example, if ‖***t****n*‖2 = {2, 8, 6, 5, 1}, then its sorted version {1, 2, 5, 6, 8} corresponds to the 10th, 30th, 50th, 70th, and 90th percentiles, respectively.
2. Strain energy (Pa· µm): 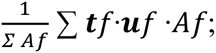; where ***t****f* and ***u****f* are the traction and displacement vector at the centroid of facet, *f*, respectively, and *Af* is the facet surface area.
3. Contractility (Pa): This metric can be defined as the mean contribution of the traction vectors towards (+) (or away from, (-)) the spheroid centroid. It is calculated as: 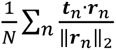 where ***r****_n_* is a unit vector that points from a node n to the spheroid centroid, · is the dot product.

### Computational Model

The model describes a spheroid domain, Ω*_s_*, embedded within a hydrogel domain, Ω*_h_*, such that the reference configuration is Ω_0_ = Ω_s_ ∪ Ω_h_. We assumed that cell contraction can result in a deformation of the domain over time *t* into some current configuration Ω*_t_*. This deformation of the domain over time is given by **x**(**X**, t) = **X** + **u**(**X**, t), where **u** is the displacement field. The deformation gradient, **F**, describes the deformation within the theory of nonlinear continuum mechanics:

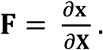

This geometric deformation tensor can be decomposed as

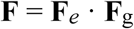

where **F***_e_* represents the elastic deformation tensor, associated with the stress response of the material, and **F***_g_* the inelastic growth tensor. The growth deformation tensor **F***_g_* can be modelled as the general form:

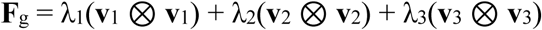

where λ_1_, λ_2_ and λ_3_ represent the contraction stretches along the orthonormal vectors **v**_1_, **v**_2_ and **v**_3_, respectively. We assume λ_i_ for *i* = 1, 2, 3 the orthonormal directions to be proportional to the ratio of cellularity:

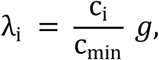

where C_i_ is the number of cells along direction *i*, C*_min_* is the minimum number of cells in one direction, and *g* represents the capacity of contraction per cell.

Simulations were performed on an idealised 380 µm cube containing a 66 µm diameter spheroid at its centre, discretised using a tetrahedral finite element mesh. Both spheroids and hydrogels were modelled as elastic materials. The Poisson’s ratio of the hydrogel was set to ν = 0.3, the same value used in the TFM analysis to reconstruct the experimental displacement fields, so that model and data rely on the same constitutive assumption. Since the hydrogels behave predominantly elastically (G’’ ≪ G’), the shear modulus can be approximated by the storage modulus, G ≈ G’, and, therefore, isotropic linear elasticity, E = 2G’(1+ ν) can be assumed. Therefore, for the soft hydrogel, we used E*_h_*= 1300 Pa, whereas for the stiff one, E*_h_* was set to 3900 Pa. The spheroid stiffness, E*_s_*, was set to 5000 Pa, higher than that of either hydrogel^21^. This choice is supported by Brillouin microscopy data, that showed the spheroid to be stiffer than the surrounding hydrogel matrix. However, the Brillouin measurements could not be used to obtain a quantitative value of E*_s_*since both the mass density and the refractive index are difficult to determine reliably, and with spatial resolution, in a heterogeneous biological tissue such as a spheroid. Spheroid contraction was prescribed as a uniform, linearly increasing deformation reaching *g* at the final state, i.e. *f_g_*(*t* = 72*h*) = *g*. Based on the experimental observations, we assumed twice as many cells along the z axis compared to the other two principal directions, resulting in λ_3_ = 2λ_1_ = 2λ_2_. Also based on the experimental displacements, we estimated *g* to be 0.35. All simulations were performed using the open-source multiphysics simulation framework 4C under quasi-static conditions, neglecting inertial effects^55^. For the study of the different parameters, QUEENS^56^ was used to schedule the simulations.

### Statistical Analysis

Data are presented as mean ± standard deviation (SD) unless stated otherwise. Statistical analyses and graph plots were performed using GraphPad Prism 10 (GraphPad Software). Normality was assessed with D’Agostino Pearson normality test. To compare more than two groups of normally distributed data, a one-way ANOVA was performed with Tukey’s post-hoc test for multiple comparisons or a Kruskal-Wallis test if normality was not met. To compare two or more groups and two conditions, a two-way ANOVA was performed with Tukey’s post-hoc test for multiple comparisons unless stated otherwise. Grubb’s test was used to identify outliers and data removal was kept to minimum. Where applicable, the number of independent experiments (N) and the number of spheroids or independent hydrogels (n) are stated. P values indicate significance *≤ 0.05, **≤ 0.01, ***≤ 0.001, ****≤ 0.0001. No statistical significance is represented by the absence of an asterisk.

## Acknowledgements

M.S-S acknowledges funds from ERC AdG Devise (101054728), ERC PoC grant (Factorink, 101248403). We acknowledge the support from EPSRC through Mechanomeds (EPSRC HT2050 grant EP/X033554/1). S.H-R would like to thank the Alexander von Humboldt Foundation and the Carl Friedrich von Siemens Foundation for their support through the Alexander von Humboldt Postdoctoral Fellowship Program. IBEC is a recipient of a Severo Ochoa Award of Excellence from MINCIN and a member of the CERCA Programme/Generalitat de Catalunya.

