## supplementary material for "Matrix mechanics gate the expression of spontaneous polarity in multicellular aggregates"

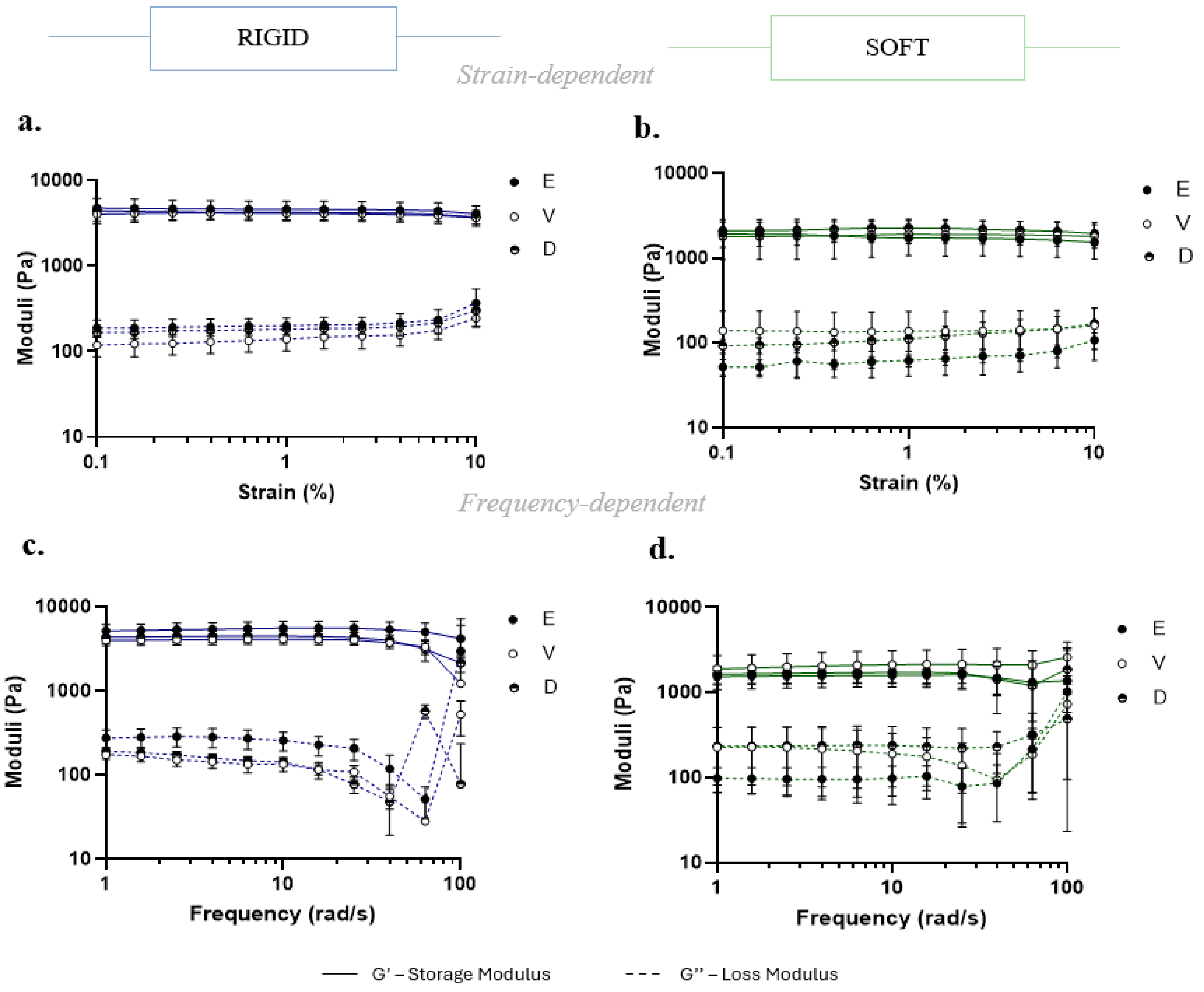


**Figure S1: Strain and frequency-dependent mechanical behaviour of FN-PEG hydrogels.**
**(a-b)** Strain-dependent behaviour of soft and rigid FN-PEG elastic, viscoelastic, and degradable hydrogels. Strain sweeps were conducted at a constant angular frequency of 10 rad/s in the range of 0.1% to 10% strain. Both storage (G'; solid lines) and loss moduli (G", dotted lines) are depicted on the graphs for all hydrogel conditions. RE: n = 9, RV: n = 8, RD: n = 7 and SE: n = 5, SV: n = 5, SD: n = 5 hydrogels **(c-d)** Frequency-dependent behaviour of soft and rigid FN-PEG hydrogels. Frequency sweeps were conducted at a constant oscillatory strain of 1% (in the case of rigid hydrogels) or 10% (in the case of soft) in the range of 1 to 100 rad/s. Both storage (G'; solid lines) and loss moduli (G", dotted lines) are depicted on the graphs for all hydrogel conditions. n = 3-6 hydrogels per experimental group. RE: n = 4, RV: n = 3, RD: n = 4 and SE: n = 5, SV: n = 6, SD: n = 5 hydrogels Data are shown as mean ± SD.

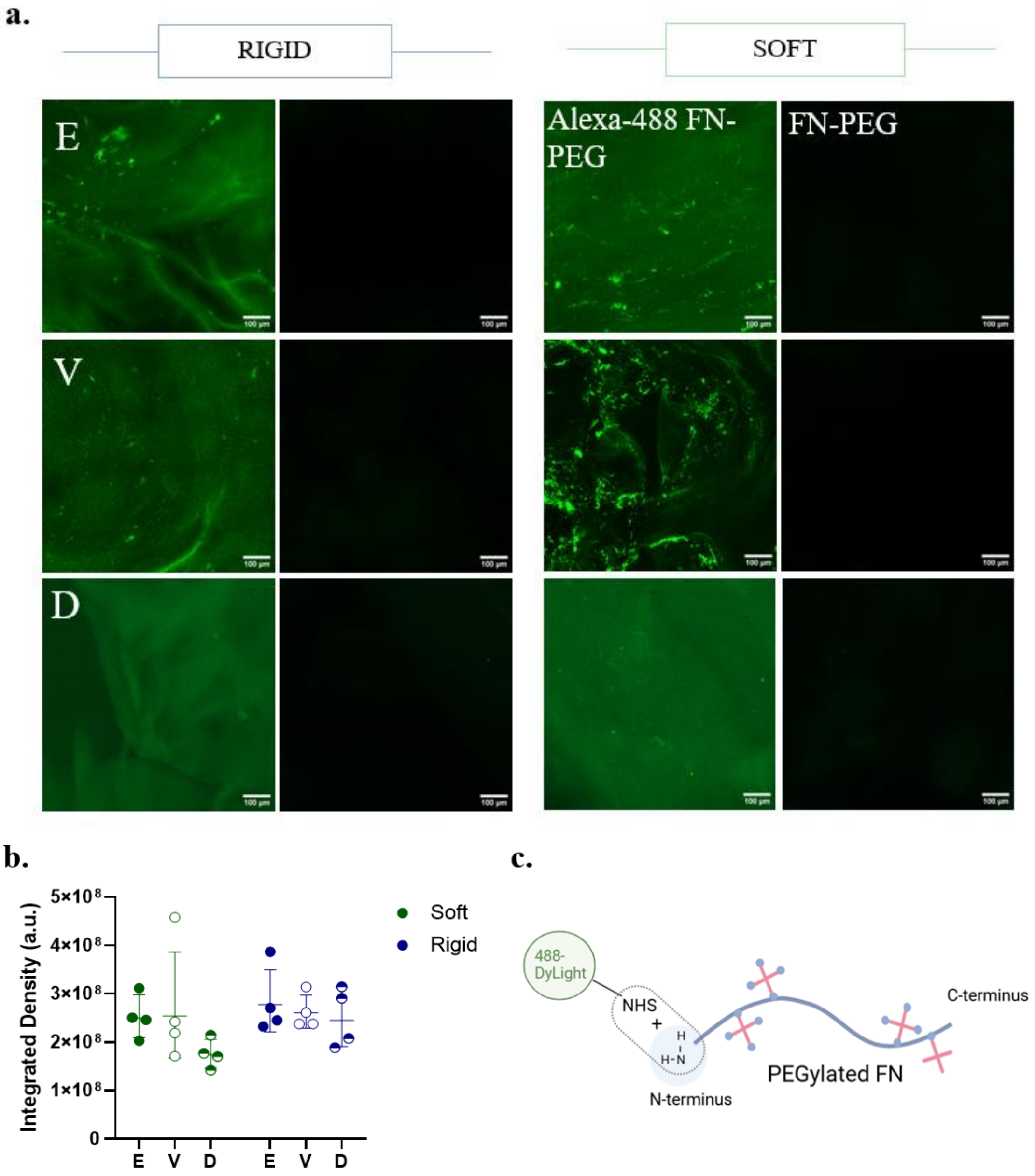


**Figure S2: Fibronectin (FN) distribution in FN-functionalised hydrogels.**
**(a)** Representative images of rigid and soft elastic (E), viscoelastic (V), and degradable (D) hydrogels with fluorescently labelled PEGylated fibronectin (left). Control hydrogels made using unlabelled PEGylated fibronectin are also shown (right) for each hydrogel type. Scale bar = 100 µm. **(b)** Quantification of resulting fibronectin intensity (integrated density, a.u.). Two images per hydrogel were acquired and three areas per image were analysed. n = 4 images analysed from 2 independent hydrogels per experimental condition. Data is shown as mean ± SD, two-way ANOVA followed by Tukey's post-hoc multiple comparisons test. **(c)** Fibronectin labelling was achieved using DyLight 488 NHS ester dye. The chemical structure of the DyLight agent comprises of an NHS ester-activated moiety that labels proteins at primary amines ($-NH_2$) existing at the N-terminus.

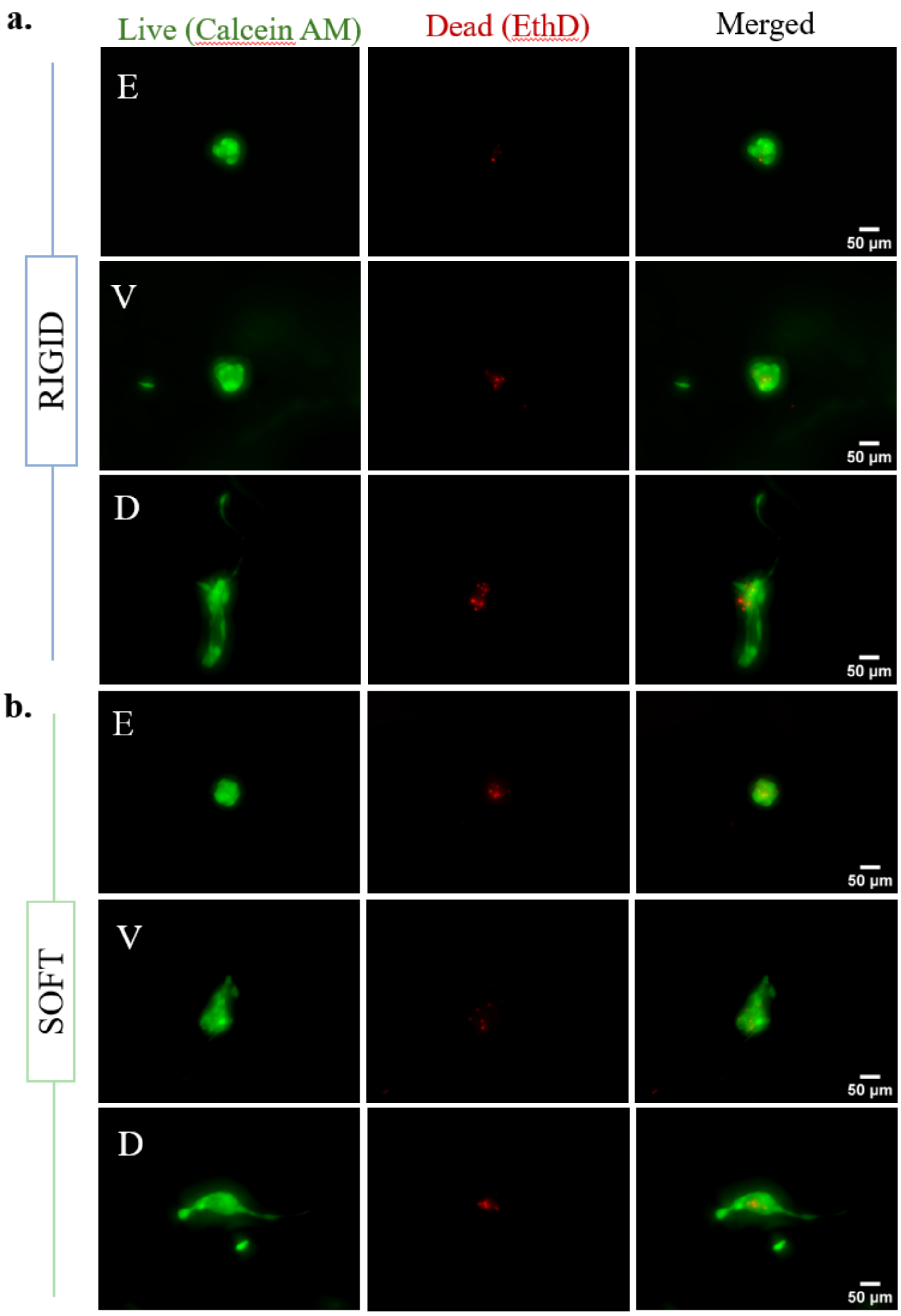


**Figure S3: Viability of hMSC spheroids encapsulated in FN-PEG hydrogels with different mechanical properties.**
**(a)** Representative images of stained live (left) and dead (middle) hMSC spheroids encapsuled in elastic, viscoelastic and degradable rigid and **(b)** soft hydrogels. Green represents live cells with normal enzymatic activity, stained by calcein AM. Red shows dead cells stained with ethidium homodimer (EThD), a nucleic acid stain that can only stain cells with compromised cell membranes. Scale bar = 50 µm.

a.

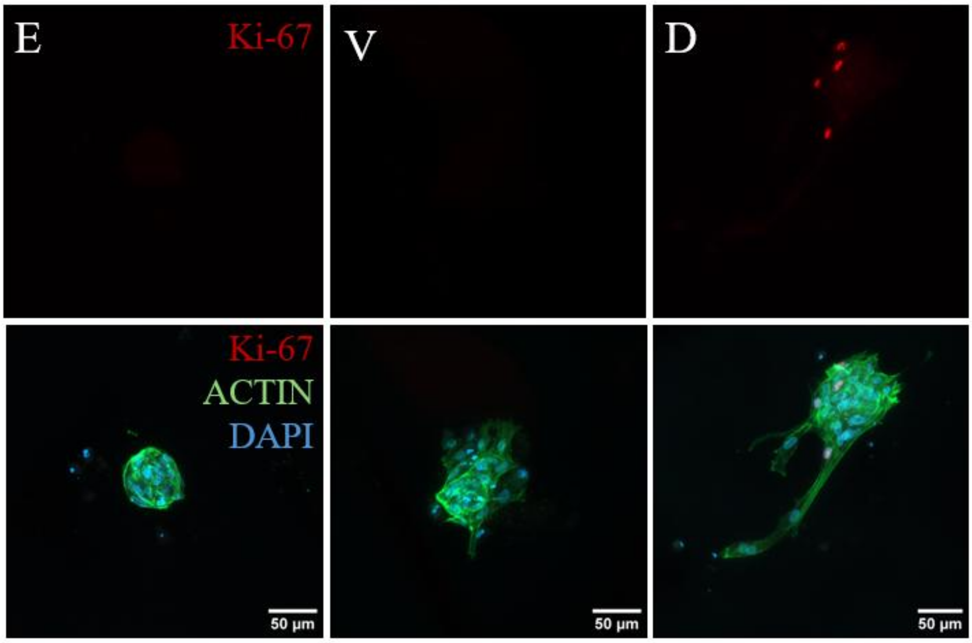


b.

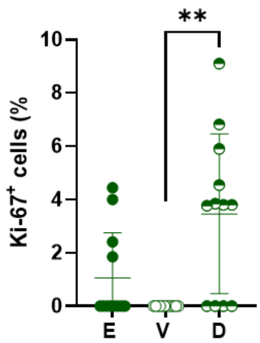


**Figure S4: Proliferation of hMSC spheroids in soft elastic, viscoelastic and degradable hydrogels.**

**(a)** Representative immunofluorescence images of Ki-67 staining (actin cytoskeleton; green, nucleus; blue, Ki-67; red) of hMSC spheroids encapsulated in soft elastic, viscoelastic, and degradable hydrogels at day 3. Scale bar = 50 µm. **(b)** Quantification of the percentage of Ki-67+ cells in spheroids in each hydrogel condition. SE: n = 12, SV: n = 12, SD: n = 12 spheroids, N = 2. Data is shown as mean ± SD, Kruskal-Wallis test. *p-value < 0.05.

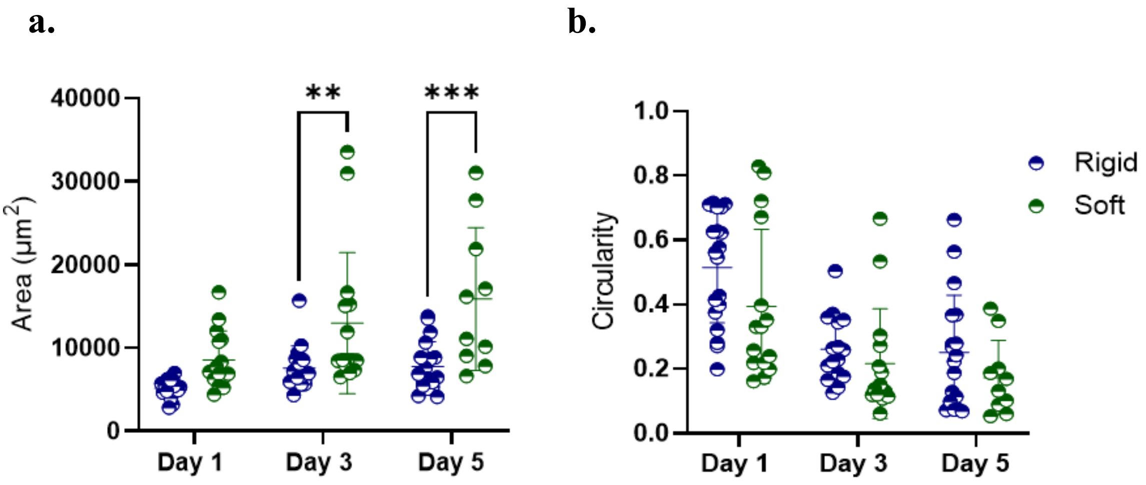


**Figure S5: Effect of hydrogel stiffness on degradation-mediated spheroid spreading**
**(a)** Quantified spheroid area and **(b)** circularity in rigid (blue) and soft (green) degradable hydrogels at respective time points. Data are shown as mean ± SD. RD: n = 16-19 and SD: n= 10-15 spheroids per condition, N = 3. Two-way ANOVA with Sidak's post-hoc test for multiple comparisons. **p-value < 0.01, ***p-value < 0.001

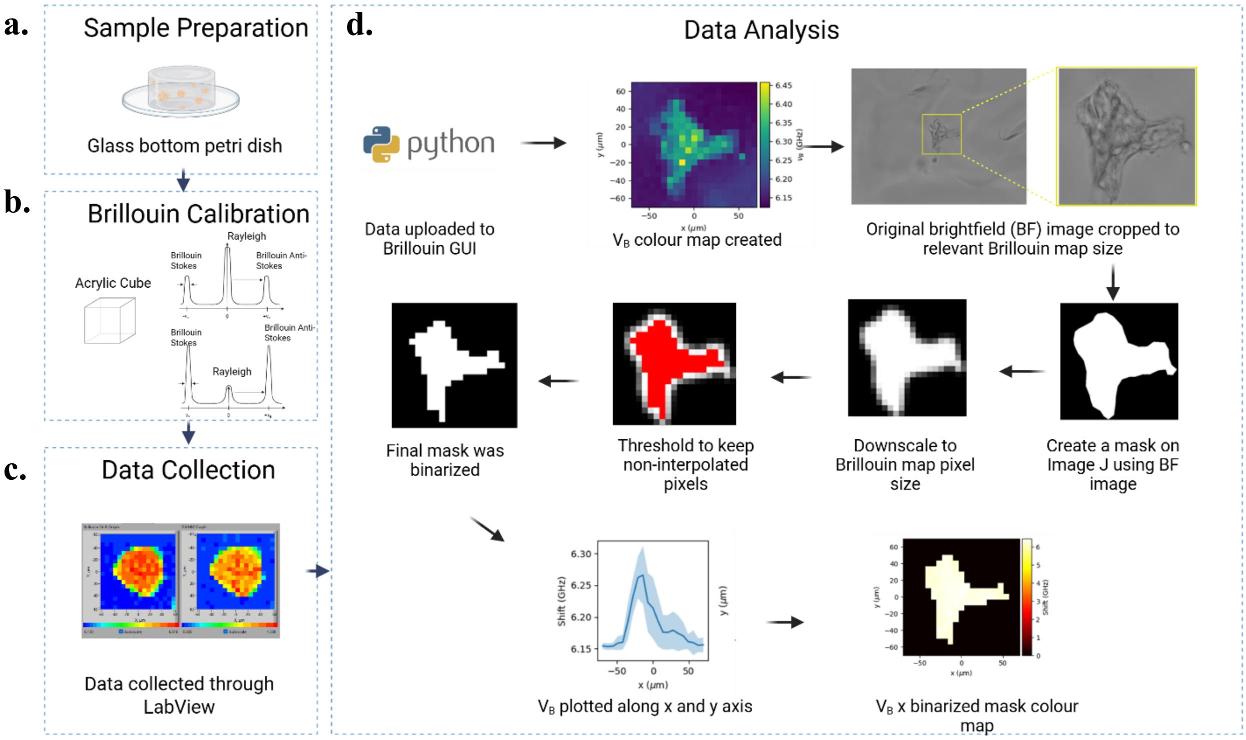


**Figure S6: Overview of Brillouin microscopy experiments, including experimental set up and analysis workflow.**

Brillouin experiments were conducted using the following step: **(a)** Sample preparation: PEG hydrogels samples with encapsulated spheroids were prepared and transferred to glass bottom petri dishes prior to scanning. **(b)** Spectra calibration: Brillouin spectra calibration was conducted to reduce Rayleigh light and increase Brillouin peak intensity. **(c)** Data collection: Three spheroids per hydrogel were measured from three technical replicates and two biological replicates. **(d)** Data analysis: For data analysis, a graphical user interface (GUI) using python was prepared in combination with ImageJ (used to create and downscale masks) to perform batch data analysis.

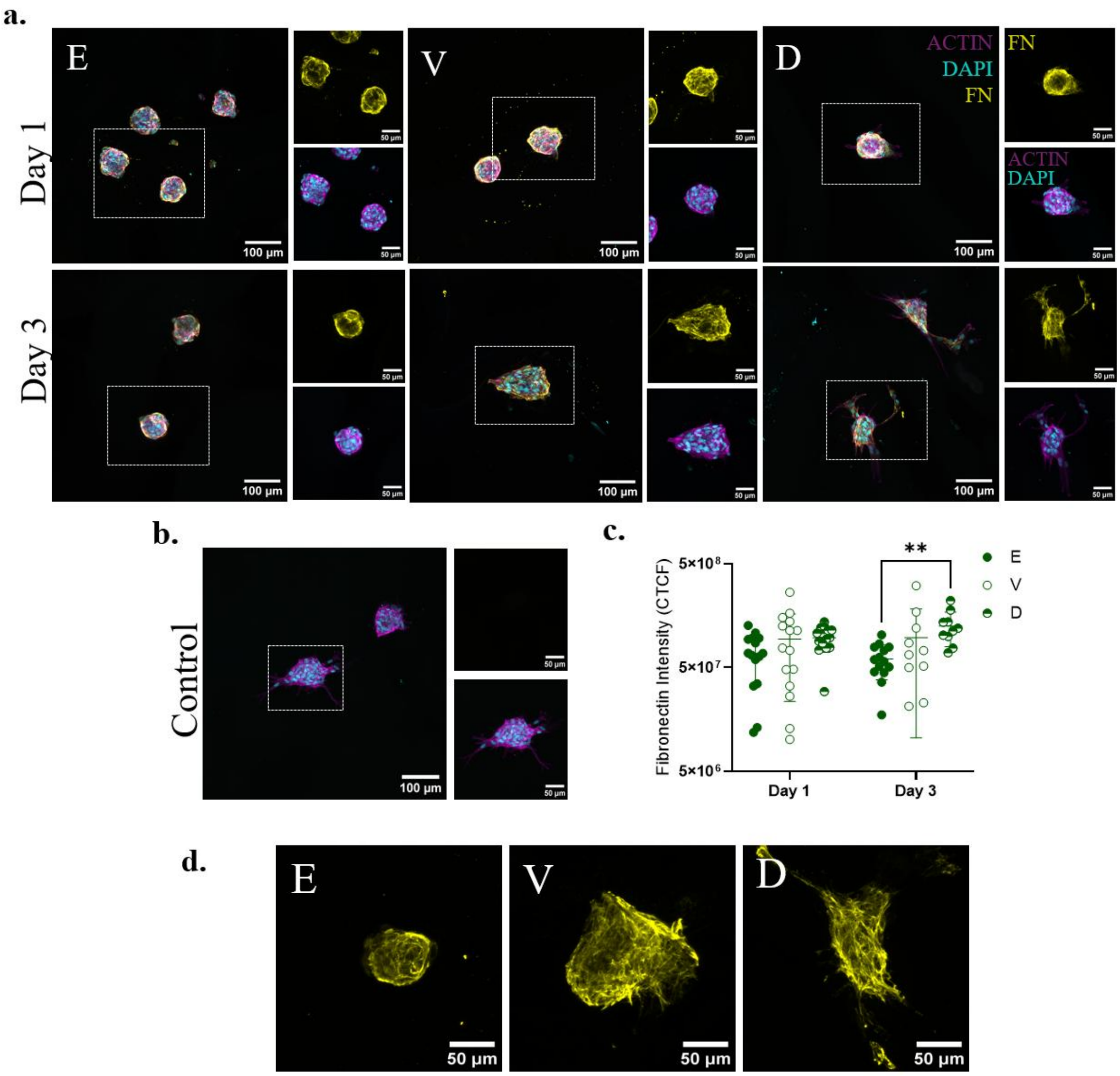


**Figure S7: Fibronectin production by hMSC spheroids in soft elastic (E), viscoelastic (V) and degradable (D) hydrogels.**

**(a)** Representative immunofluorescence images of cellular-secreted fibronectin (FN) production (actin cytoskeleton; magenta, nucleus; cyan, fibronectin; yellow) in hMSC spheroids encapsulated in SE, SV, and SD hydrogels at days 1 and 3 (top and bottom panel, respectively). **(b)** Representative image of secondary-only control (at day 3), showing no FN signal when stained only with secondary antibody. Scale bar = 100 µm in main image panels and scale bar = 50 µm in insets. **(c)** Fibronectin fluorescence intensity quantification using corrected total cell fluorescence (CTCF) analysis. Results are presented as mean ± SD. SE: n = 14-15, SV: n = 10-15 SD: n = 10-15 spheroids, N = 2. Two-way ANOVA with Tukey's post hoc test for multiple comparisons, **p-value < 0.01. **(d)** Magnified immunofluorescence images showing FN distribution and organisation in SE, SV, and SD hydrogels on day 3.

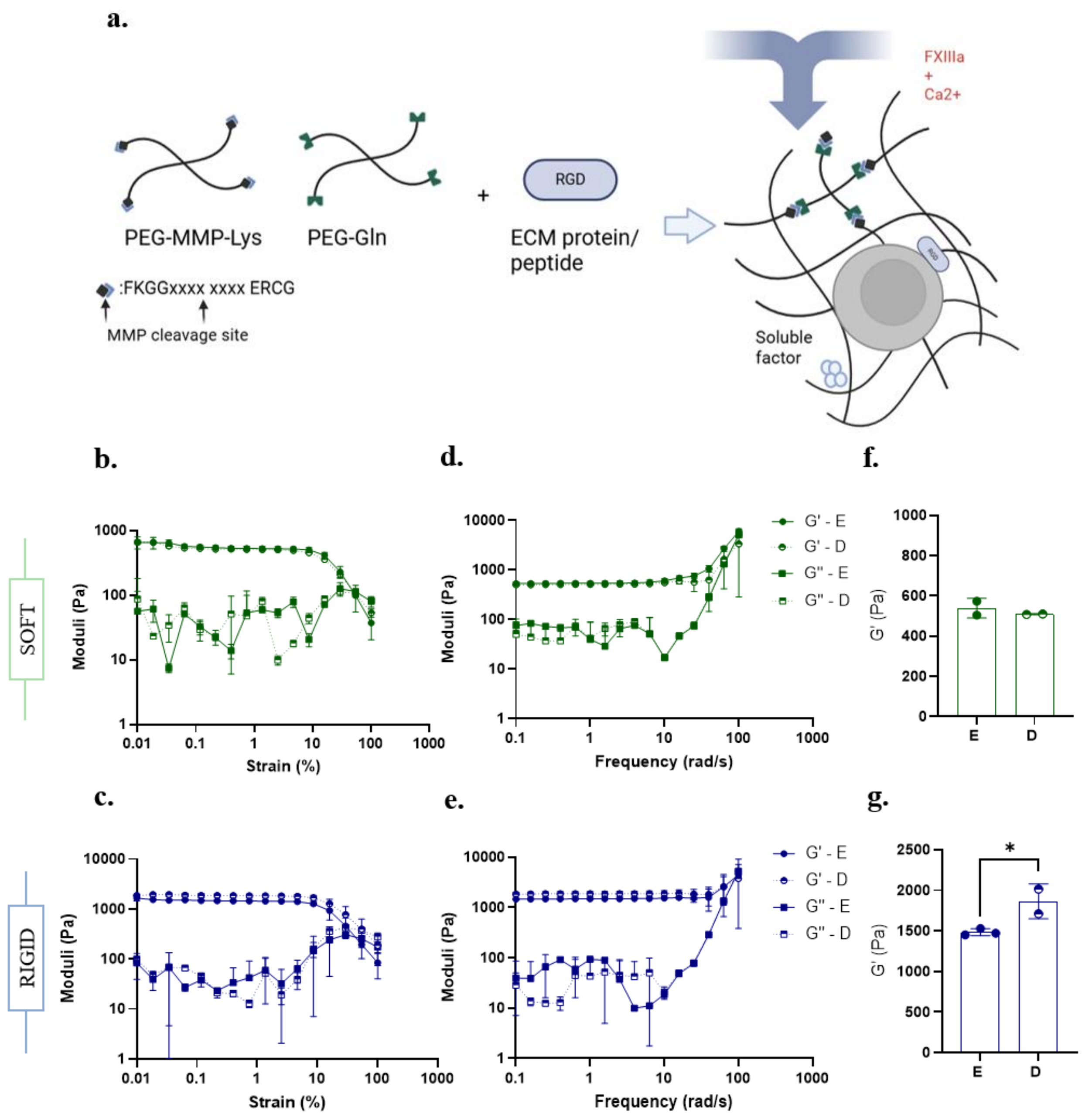


**Figure S8: Mechanical characterisation of PEG-VS hydrogels containing fluorescent beads.** **(a)** Schematic representation of PEG-VS hydrogel formation. Image adapted from Ranga et al. 2014. **(b-c)** Strain-dependent and **(d-e)** frequency-dependent behaviour of soft (green) and rigid (blue) elastic (non-degradable) and degradable RGD-functionalised PEG-VS hydrogels, respectively. Strain sweeps were conducted at an angular frequency of 1 rad/s, with strain amplitudes ranging from 0.01 to 100%, demonstrating the linear viscoelastic regime (LVE) of each hydrogel composition. Frequency sweeps were performed with a constant strain of 1% while the angular frequency increased from 0.1 to 100 rad/s. **(f-g)** The storage modulus (G') of soft and rigid elastic and degradable PEG-VS hydrogels reported at a frequency of 1 rad/s. SE: n = 2, SD: n = 2 and RE: n = 3, RD: n = 2 hydrogels. Data are shown as mean ± SD, unpaired two-tailed t-test, *p-value < 0.05.

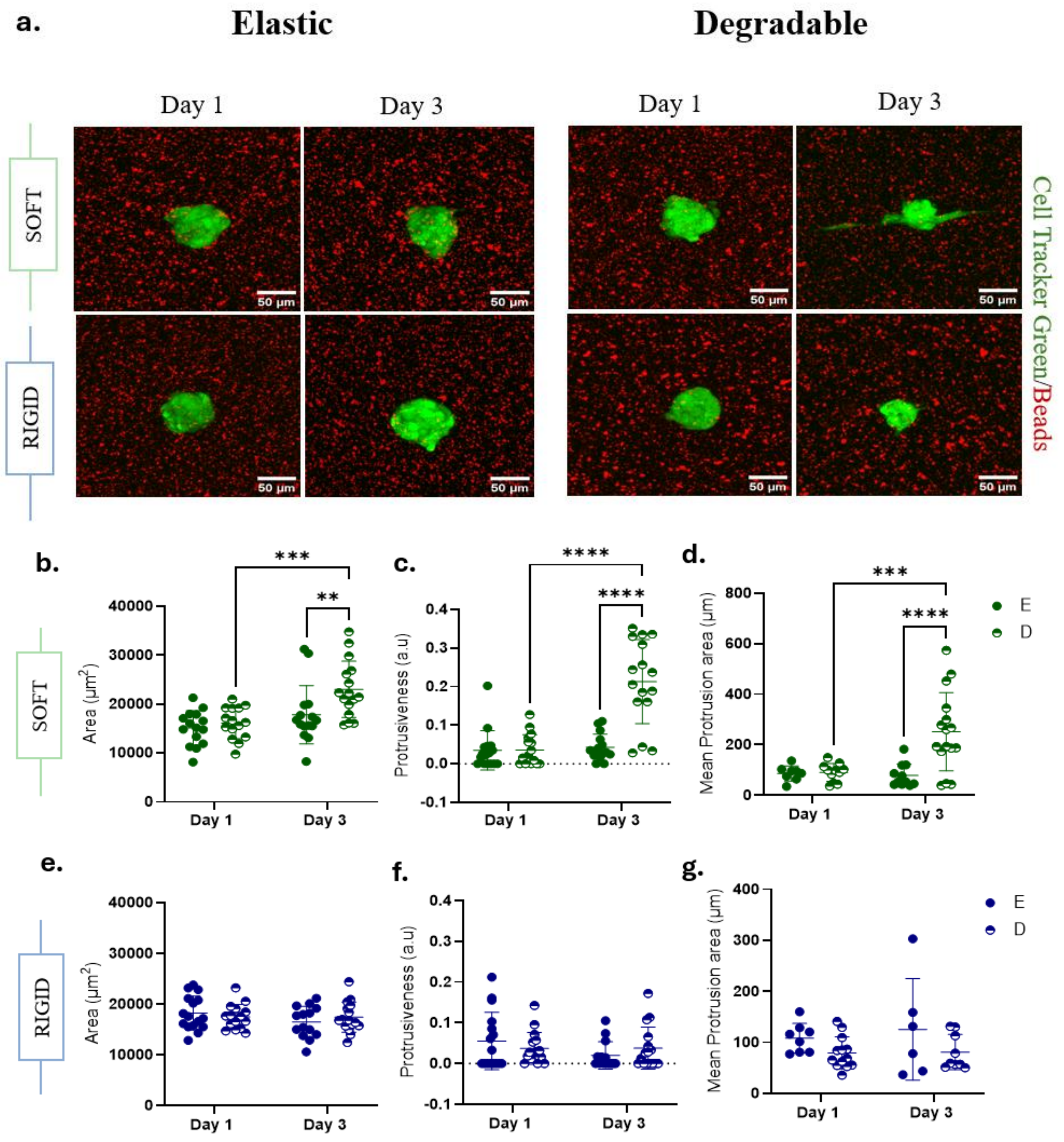


**Figure S9: hMSC spheroid morphology in soft and rigid elastic (E) and degradable (D) hydrogels. (a)** Representative maximally projected images of beads (in red) and cell-tracker green-labelled hMSC spheroids (in green) for elastic and degradable soft and rigid hydrogels at days 1 and 3. Scale bars = 50 µm. **(b-d)** Quantification of the spheroid area, protrusiveness and mean protrusion area of spheroids in elastic and degradable soft and **(e-g)** rigid RGD-functionalised PEG-VS hydrogels. SE: n = 15-16, SD: n = 16 and RE: n = 14 – 16, RD: n = 16 spheroids, N = 2. Data are shown as mean ± SD, Two-way ANOVA with Sidak's post hoc test for multiple comparisons, *p-value < 0.05, ***p-value < 0.001, ****p-value < 0.0001.

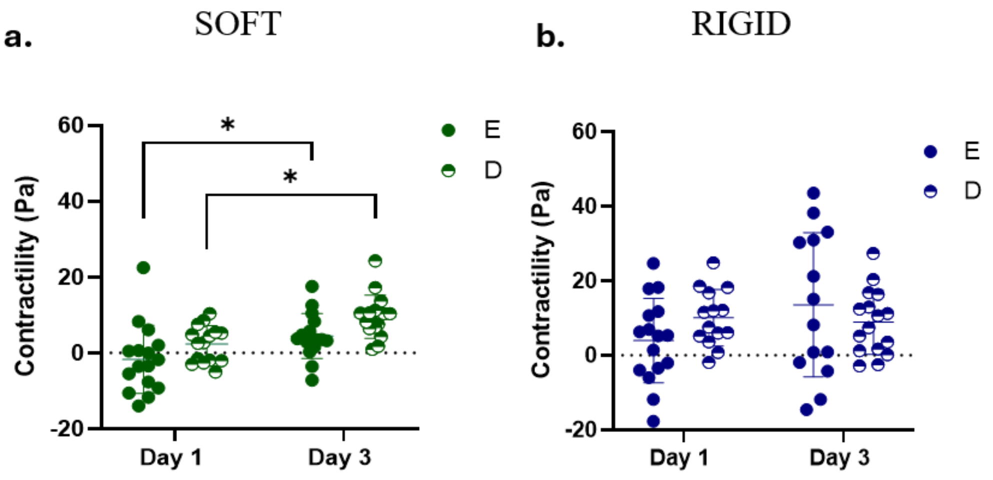


**Figure S10: Contractility of hMSC spheroids in soft and rigid elastic (E) and degradable (D) over time.**
**(a)** Contractility, defined as the mean contribution of the traction vectors towards (+; pulling) or away (-; pushing) of the spheroid centre of hMSC spheroids in soft and **(b)** rigid hydrogels at respective time points. Data are shown as mean ± SD. SE: n = 15-16, SD: n = 16 and RE: n = 14 – 16, RD: n = 16 spheroids, N = 2. Two-way ANOVA with Tukey's post-hoc test for multiple comparisons, *p-value < 0.05.

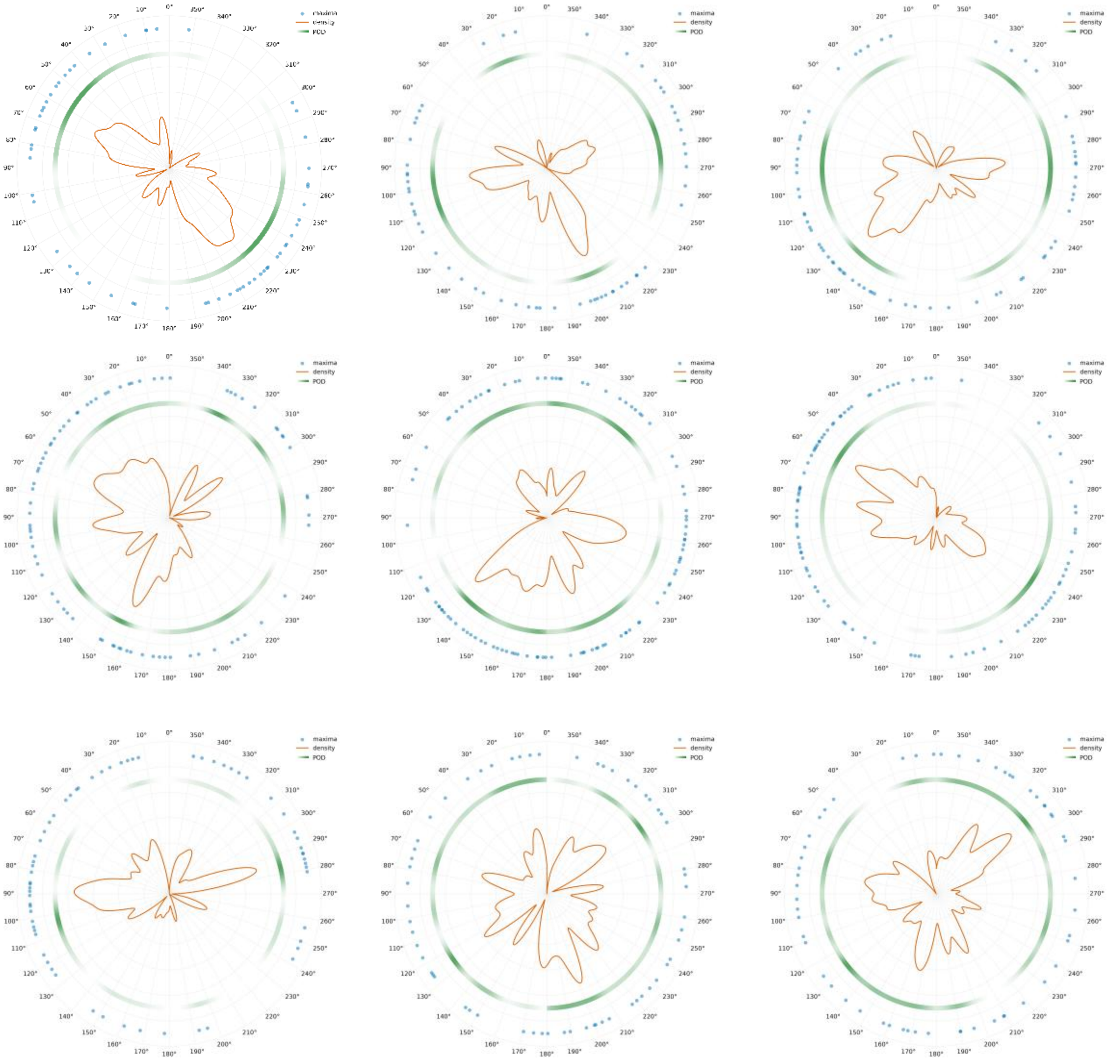

**Figure S11: Spatial organisation of paxillin around the rim of hMSC spheroids in soft elastic hydrogels.**

Polar plots showing the angular distribution of paxillin maxima (blue dots), density of maxima (orange line) and product of opposite density (POD; green line) for individual spheroids encapsulated in soft elastic hydrogels.

### Appendix - Computational model

#### 1.1 Influence of the Poisson's ratio

The Poisson's ratio of the hydrogels was not measured experimentally, so we treated ν as an uncertain parameter (uniform in [0.30, 0.48], n = 30 per hydrogel) and propagated it through the model. Because the model is driven kinematically, by a prescribed contraction of a spheroid that is stiffer than the surrounding gel, the hydrogel displacements are almost insensitive to ν. The 99$^{th}$ percentile displacement varied by less than 0.5% (coefficient of variation 0.27% for the soft and 0.41% for the stiff hydrogel), well below the experimental resolution. Since displacements are the only quantity measured and used for calibration, the calibrated parameters are robust to the uncertainty in ν. Stresses in contrast, do depend on ν. In the soft hydrogel, ν = 0.48 produces clearly higher Cauchy stresses than ν = 0.30, and they reach further into the gel, because the bulk modulus K = E/[3(1−2ν)] rises steeply as ν approaches 0.5. Derived quantities such as stresses and tractions are therefore conditional on the assumed ν, as they are in conventional traction force microscopy.

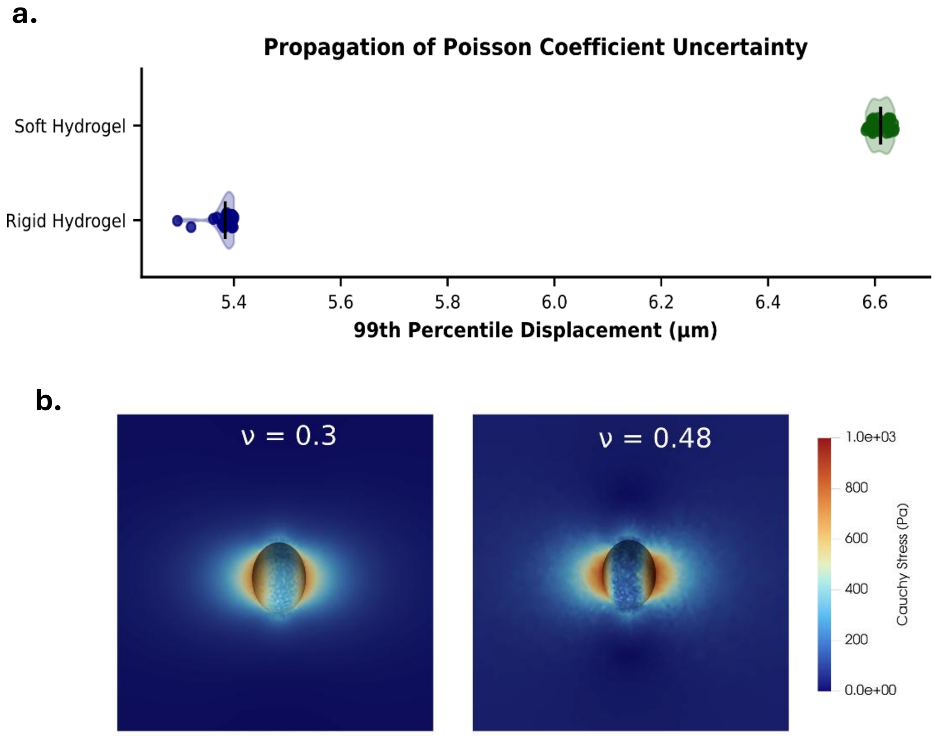


**Figure S12: Sensitivity of the predicted displacement to the assumed material properties.**
**(a)** 99th percentile of the hydrogel displacement magnitude for the soft and rigid hydrogels across n = 30 realisations of ν ∈ [0.30, 0.48]. Dots are individual simulations; black lines mark the mean. **(b)** Cauchy stress in the soft hydrogel at the end of the simulation for ν = 0.30 (left) and ν = 0.48 (right).

#### 1.2 Uncertainty of the spheroid stiffness

The Young's modulus, $E_s$, of the spheroid in situ was hard to obtain experimentally from Brillouin frequency shift values. As with the Poisson's ratio (Section 1.1), we therefore checked how sensitive the model is to this parameter. We varied $E_s$ (255–9700 Pa) and the contraction rate $g$ (−0.75 to −0.16) at the same time over 50 simulations and computed the 99$^{th}$ percentile of the displacement magnitude in the matrix. The results suggest that, for this model, the displacements depend mainly on contractility. They increased clearly with |g| (Pearson's r = 0.92), whereas changing Es by almost two orders of magnitude had little effect (r = 0.14). Simulations that matched the experimental range used intermediate contraction rates (|g| ≈ 0.41–0.55) and stiffnesses across nearly the whole sampled range. The uncertainty in the spheroid stiffness therefore seems to have a limited impact on the calibration of the contraction rate.

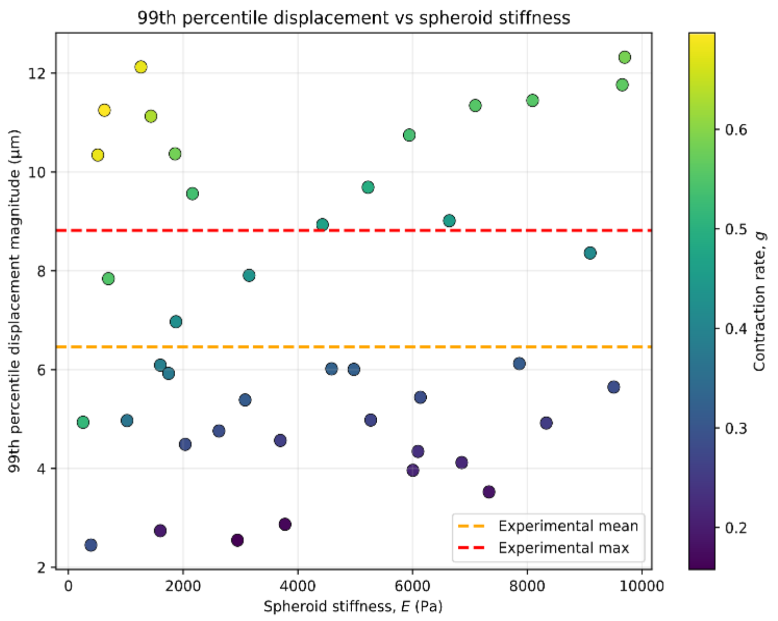


**Figure S12: Combinations of spheroid stiffness and contractility that can predict generated experimental displacements.**
Young's modulus of the spheroid ranges from 100Pa to 10kPa. The plot shows the 99$^{th}$ percentile of the displacement obtained for a specific Es, coloured by the contractility *g*.

**Table S1**: Hydrogel formulations (for 50 µl hydrogels).

| Reagents | SE | SV | SD | RE | RV | RD | Stock concentrations (mg/ml) |
|---|---|---|---|---|---|---|---|
| **FN-PEG (µl)** | 10.00 | 10.00 | 10.00 | 10.00 | 10.00 | 10.00 | 2.50 |
| **PEG-Mal (µl)** | 4.00 | 4.00 | 4.00 | 5.00 | 8.00 | 5.00 | 500.00 |
| **HS-PEG-SH (µl)** | 2.00 | 20.00 | 1.00 | 10.00* | 8.00* | 5.00* | 400.00 / 200.00* |
| **VPM (µl)** | - | - | 1.70 | - | - | 4.25 | 200.00 |
| **PBS** | 34.00 | 17.00 | 33.30 | 25.00 | 24.00 | 25.75 | N/A |
| **Total Volume (µl)** | **50.00** | **50.00** | **50.00** | **50.00** | **50.00** | **50.00** | **N/A** |
| **% PEG-Mal (w/v)** | **4.00%** | **4.00%** | **4.00%** | **5.00%** | **8.00%** | **5.00%** | **N/A** |

**Table S2:** Statistically significant differences in hMSC spheroid area ($\mu m^2$) and circularity over time in *rigid* elastic (E), viscoelastic (V) and degradable (D) hydrogels. *p < 0.05, **p < 0.01 ***p < 0.001, ****p <0.0001.

| Statistically significant differences in morphological parameters over time | | | | |
|---|---|---|---|---|
| | Area | | Circularity | |
| **RE:** Day 1 vs Day 3 | ns | 0.8683 | ns | 0.7059 |
| **RE:** Day 1 vs Day 5 | * | 0.0196 | ns | 0.5750 |
| **RE:** Day 3 vs Day 5 | ns | 0.0730 | ns | 0.9386 |
| **RV:** Day 1 vs Day 3 | ns | 0.1020 | ns | 0.6290 |
| **RV:** Day 1 vs Day 5 | ns | 0.8010 | ns | 0.3960 |
| **RV:** Day 3 vs Day 5 | ns | 0.4855 | ns | 0.1048 |
| **RD:** Day 1 vs Day 3 | ** | 0.0022 | **** | <0.0001 |
| **RD:** Day 1 vs Day 5 | *** | 0.0007 | **** | <0.0001 |
| **RD:** Day 3 vs Day 5 | ns | 0.9744 | ns | 0.9818 |

**Table S3:** Statistically significant differences in hMSC spheroid area ($\mu m^2$) and circularity over time in *soft* elastic (E), viscoelastic (V) and degradable (D) hydrogels. *p < 0.05, **p < 0.01 ***p < 0.001, ****p <0.0001.

| Statistically significant differences in morphological parameters over time | | | | |
|---|---|---|---|---|
| | Area | | Circularity | |
| **SE:** Day 1 vs Day 3 | ns | 0.8647 | ns | 0.3896 |
| **SE:** Day 1 vs Day 5 | ns | 0.8457 | * | 0.0397 |

| | | | | |
|---|---|---|---|---|
| **SE:** Day 3 vs Day 5 | ns | 0.9967 | ns | 0.4386 |
| **SV:** Day 1 vs Day 3 | ns | 0.4983 | ns | 0.7379 |
| **SV:** Day 1 vs Day 5 | **** | <0.0001 | ns | 0.8865 |
| **SV:** Day 3 vs Day 5 | **** | <0.0001 | ns | 0.4467 |
| **SD:** Day 1 vs Day 3 | ** | 0.0088 | ** | 0.0057 |
| **SD:** Day 1 vs Day 5 | **** | <0.0001 | ** | 0.0017 |
| **SD:** Day 3 vs Day 5 | ns | 0.1904 | ns | 0.7692 |

**Table S4:** Statistically significant differences in hMSC spheroid area ($\mu m^2$) and circularity between *soft* elastic (E), viscoelastic (V) and degradable (D) PEG and FN-PEG hydrogels at day 1. *p < 0.05, **p < 0.01 ***p < 0.001, ****p <0.0001.

| **Statistically significant differences in morphological parameters between SE, SV and SD PEG and FN-PEG hydrogels** | | | | |
|---|---|---|---|---|
| | **Area** | | **Circularity** | |
| **PEG:** SE vs SV | * | 0.0304 | ns | 0.0938 |
| **PEG:** SE vs SD | ** | 0.0071 | **** | <0.0001 |
| **PEG:** SV vs SD | ns | 0.9714 | * | 0.0255 |
| **FN-PEG:** SE vs SV | **** | <0.0001 | **** | <0.0001 |
| **FN-PEG:** SE vs SD | ** | 0.0064 | **** | <0.0001 |
| **FN-PEG:** SV vs SD | ns | 0.0666 | ns | 0.6055 |

**Table S5:** Statistically significant differences in displacements (μm) and traction forces (Pa) between rigid and soft elastic (E) and degradable (D) hydrogels on day 1 and 3. *p < 0.05, **p < 0.01 ***p < 0.001, ****p <0.0001.

| **Statistically significant differences in morphological parameters over time** | | | | |
|---|---|---|---|---|
| | **Displacements** | | **Traction Forces** | |
| **Day 1:** SE vs RE | **** | <0.0001 | ns | 0.6548 |
| **Day 1:** SD vs RD | ** | 0.0026 | *** | 0.0001 |
| **Day 3:** SE vs RE | ** | 0.0012 | **** | <0.0001 |
| **Day 3:** SD vs RD | **** | <0.0001 | **** | <0.0001 |
